# Integrator terminates promoter-proximal Pol II to generate *C. elegans* piRNA precursors

**DOI:** 10.1101/2020.05.01.072298

**Authors:** Toni Beltran, Elena Pahita, Subhanita Ghosh, Boris Lenhard, Peter Sarkies

## Abstract

Piwi-interacting RNAs (piRNAs) play key roles in germline development and genome defence in metazoans. In *C. elegans*, piRNAs are transcribed from >15000 discrete genomic loci by RNA polymerase II, resulting in 28 nt short-capped piRNA precursors. Here we investigate transcription termination at piRNA loci. We show that the Integrator complex, which terminates snRNA transcription, is recruited to piRNA loci. We show that the catalytic activity of Integrator cleaves nascent capped piRNA precursors associated with promoter-proximal Pol II, resulting in termination of transcription. Loss of Integrator activity, however, does not result in transcriptional readthrough at the majority of piRNA loci. Our results draw new parallels between snRNA and piRNA biogenesis in nematodes, and provide evidence of a role for the Integrator complex as a terminator of promoter-proximal RNA polymerase II.

**Highlights:**

- Integrator localises to sites of piRNA biogenesis in nematodes
- Integrator cleaves nascent RNAs associated with promoter-proximal Pol II at piRNA loci to release short capped piRNA precursors from chromatin
- Repression of Pol II elongation at the majority of piRNA loci is independent of Integrator

## Introduction

Piwi-interacting RNAs (piRNAs) are a specific class of small RNAs essential for germline development and transposable element silencing in metazoans (Ozata et al., 2019). piRNA biogenesis is best characterized in the fruit fly *Drosophila melanogaster* and in the nematode *Caenorhabditis elegans*. In *D. melanogaster*, piRNAs are transcribed from a small number of genomic clusters to produce >10 kb long piRNA precursors. piRNA precursors are then processed by nuclease activities into 27-35 nucleotide long primary piRNAs. Upon detection of transposable element transcripts, piRNAs initiate a cycle of coupled cleavage reactions involving transposable element mRNAs and piRNA precursor RNAs. This coupling is known as the Ping-pong cycle, which allows amplification of piRNA populations while simultaneously degrading TE mRNAs (Brennecke et al., 2007; Gunawardane et al., 2007).

*C. elegans* piRNAs present remarkable differences to the *Drosophila* system. Mature piRNAs in *C. elegans* are 21 nucleotides long with a strong bias towards uracil as the first nucleotide (21U-RNAs; (Batista et al., 2008; Das et al., 2008; Ruby et al., 2006)). *C. elegans* piRNA loci are individual transcriptional units. The majority of piRNA loci localize to two ∼3 Mb cluster regions on chromosome IV containing more than 15,000 loci, and are demarcated by a conserved GTTTC motif at their promoter regions (Billi et al., 2013; Ruby et al., 2006); these loci are known as motif-dependent piRNAs or type I piRNAs. Another set of low-abundance piRNAs are generated from almost 10,000 loci distributed across the *C. elegans* genome and independently of an upstream motif; these are known as motif-independent piRNAs (Gu et al., 2012) or type II piRNAs. Both types of loci are transcribed as short 27-40 nt capped RNA precursors by RNA polymerase II (Pol II) (Gu et al., 2012). These short precursors have been proposed to result from pausing of Pol II at piRNA promoters (Beltran et al., 2019; Gu et al., 2012). We previously described a region of low melting temperature (high AT content) centered 33 nt downstream of piRNA TSSs across multiple nematode species, which may have a role in RNA polymerase II termination (Beltran et al., 2019). However, the exact mechanisms controlling this process are largely unknown. In particular, the role of Pol II-associated factors in promoting pausing and premature termination of Pol II remain unexplored.

Multiple lines of evidence suggest that the mechanisms of piRNA transcription in nematodes evolved by co-option of factors controlling the transcription of small nuclear RNAs (snRNAs). First, transcription of piRNAs requires the small nuclear RNA activating protein complex SNAPc (Kasper et al., 2014). SNAPc forms a distinct complex in the germline with the nematode-specific pseudokinase PRDE-1 (Kasper et al., 2014; Weick et al., 2014; Weng et al., 2019), which binds to the upstream Ruby motif directly (Weng et al., 2019). In addition, the GTTTC motif is found in snRNA promoters in basal nematodes (Beltran et al., 2019). Transport of piRNA precursors to their processing sites in perinuclear phase-separated P-granules is achieved by a specialised protein complex named PETISCO/PICS (Cordeiro Rodrigues et al., 2019; Zeng et al., 2019). Interestingly, components of this complex are required for the biogenesis of SL1 RNAs, the trans-splicing leaders preceding the majority of *C*.*elegans* mRNAs, which are also related to snRNAs (Cordeiro Rodrigues et al., 2019). Altogether, these observations suggest that snRNAs and piRNAs share mechanisms of transcription and processing.

In metazoans, termination of snRNA genes requires the Integrator complex (Baillat et al., 2005). Integrator is a protein complex containing 12-14 subunits, which associates co-transcriptionally with the carboxy terminal domain (CTD) of Pol II. Two of the subunits of the complex, Ints-9 and Ints-11, are homologs of CPSF73, the catalytically active subunit of the cleavage and polyadenylation complex (CPSF) responsible for the formation of 3’ ends of protein-coding gene mRNAs. Ints-11 possesses the endonuclease activity responsible for cleavage of nascent snRNAs to generate snRNA 3’ ends. This cleavage is coupled to Pol II termination.

Whilst the best described function for Integrator is at snRNA loci, Integrator may also have functions in transcription of other Pol II-dependent transcripts. Integrator was shown to promote early termination of Pol II at the HIV LTR promoter, attenuating HIV transcription (Stadelmayer et al., 2014). A role for Integrator in conferring processivity to Pol II during the elongation stage of transcription has also been reported (Gardini et al., 2014; Stadelmayer et al., 2014). Additionally, Integrator mediates the biogenesis of enhancer RNAs (eRNAs) by promoting early termination of Pol II (Lai et al., 2015). Finally, Integrator has been proposed to terminate lncRNA transcription (Nojima et al., 2018).

Here, we discover that the Integrator complex is involved in piRNA biogenesis in nematodes. We show that Integrator activity is required for accumulation of mature piRNAs and their silencing activity. Integrator associates with piRNA clusters in germ cells, and its catalytic activity is essential to produce short capped RNA precursors via co-transcriptional cleavage of nascent piRNA precursors. Our results provide new insights into the mechanisms of piRNA biogenesis in nematodes, and uncover novel functions of the Integrator complex in the regulation of promoter-proximal Pol II pausing and the biogenesis of non-coding RNAs.

## Results

### Integrator is required for the biogenesis of short piRNA precursors

Prompted by the evolutionary relationship between piRNAs and snRNAs (Beltran et al., 2019), we sought to understand whether Integrator is involved in piRNA biogenesis in *C. elegans*. We performed RNAi against the catalytic subunit of the complex, *ints-11*, and measured changes in piRNA abundance upon knockdown by sequencing small RNAs. We observed a decrease in piRNA abundance in *ints-11*-depleted worms compared to the empty vector controls (Figure 1A, Supp. Fig. 1A-C). To further test this, we assessed the silencing activity of piRNAs using a strain carrying a germline histone H2B::GFP transgene as a sensor of piRNA activity (Bagijn et al., 2012a). Upon knockdown of *ints-11*, we observed an increase in the proportion of GFP-positive animals (Supp. Fig. 1D), suggesting that Integrator functions in piRNA biogenesis.

**Figure 1.**
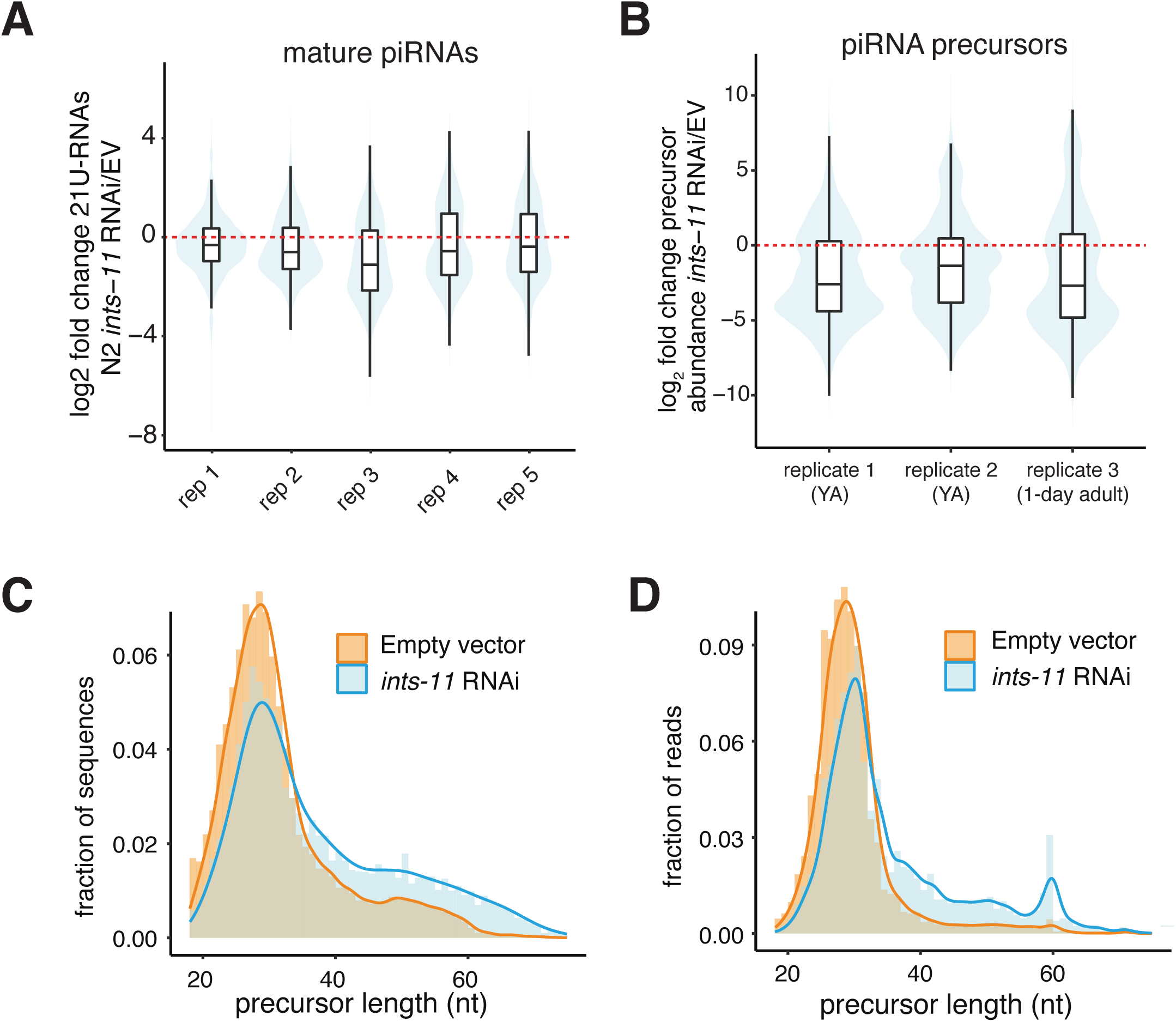
Integrator is required for motif-dependent piRNA precursor abundance and their short length. A. Distributions of log_2_ fold change in piRNA abundance in *ints-11* RNAi-treated nematodes compared to empty vector-treated nematodes. B. Distributions of log_2_ fold change in piRNA precursor abundance in *ints-11* RNAi-treated nematodes compared to empty vector-treated nematodes. C-D. Distributions of piRNA precursor length in *ints-11* RNAi-treated nematodes compared to empty vector-treated nematodes, obtained from deeply sequencing one pair of libraries (inserts sequenced up to 75nt). Distributions of total sequences (C) and total read counts (D) are shown.

To test whether Integrator promotes piRNA transcription, we performed high-throughput sequencing of short (18-36 nt) RNAs with a 5′ cap, allowing us to capture short-capped piRNA precursors. Upon knockdown of *ints-11*, we observed a decrease in piRNA precursors from piRNA loci with the characteristic GTTTC core (Ruby) motif (motif-dependent piRNAs; Figure 1B, Supp. Fig. 1E-F). We also observed a marked increase in motif-dependent piRNA precursor length upon Integrator depletion (Supp. Fig. 1G), suggesting that Integrator plays a role in termination of Pol II at motif-dependent piRNA promoters. To characterize the length distribution of piRNA precursors in greater detail, we performed high-throughput sequencing of capped RNAs up to a length of 75 nt. The previously described peak of ∼28 nt (Weick et al., 2014) was accompanied by an additional population of longer precursors up to ∼75 nt long (Figure 1C-D). This population was rare in wild type nematodes, but accumulated substantially upon *ints-11* depletion (Figure 1C-D). Altogether, these data strongly suggest a role for Integrator in the termination of short motif-dependent piRNA precursor transcripts. Interestingly, we did not detect a decrease in abundance of motif-independent piRNA precursors, suggesting that their biogenesis does not depend on Integrator (Supp. Fig. 1H).

### Integrator localizes to sites of piRNA biogenesis in germ cells

We investigated whether Integrator is recruited to sites of piRNA biogenesis. We used a strain where an endogenous deletion (*tm1616*) in the *ints-6* Integrator subunit, also known as *dic-1* in *C. elegans*, is rescued by an *ints-6*::3xFLAG::eGFP fusion transgene (Gómez-Orte et. al., 2019). INTS-6::eGFP formed discrete foci in germ nuclei (Figure 2A), which start to form in the mitotic and transition zone regions of the germline, and become most apparent in early-mid pachytene. To test whether these foci correspond to piRNA cluster regions, we crossed this strain with one carrying an mCherry-tagged version of the piRNA biogenesis factor PRDE-1, which is known to bind to piRNA clusters (Weick et al., 2014; Weng et al., 2019). The majority of mCherry::PRDE-1 foci co-localized with INTS-6::eGFP foci across the germline (Figure 2B, quantification in Supp. Fig. 2A). We observed a second population of INTS-6 foci appearing as cells progress into pachytene (Figure 2A), with mid-pachytene cells showing two clear INTS-6 foci, one of which co-localizes with PRDE-1 (Figure 2B, panels 2-3). Towards the end of pachytene, PRDE-1 foci became more diffuse and start to disappear, and INTS-6 signal in PRDE-1 regions was clearly reduced, resulting in no co-localizing foci (Figure 2B, panel 4). Interestingly, however, a single INTS-6 focus remained in late-pachytene cells well after the disappearance of PRDE-1 foci (Figure 2B, panel 5). In order to further confirm these observations, we used CRISPR-mediated genome editing to tag the endogenous INTS-11 protein with an N-terminal FLAG tag, and tested its localization by immunofluorescent staining. FLAG::INTS-11 also co-localized with PRDE-1 foci in germ cells (Supp. Fig. 2B).

**Figure 2.**
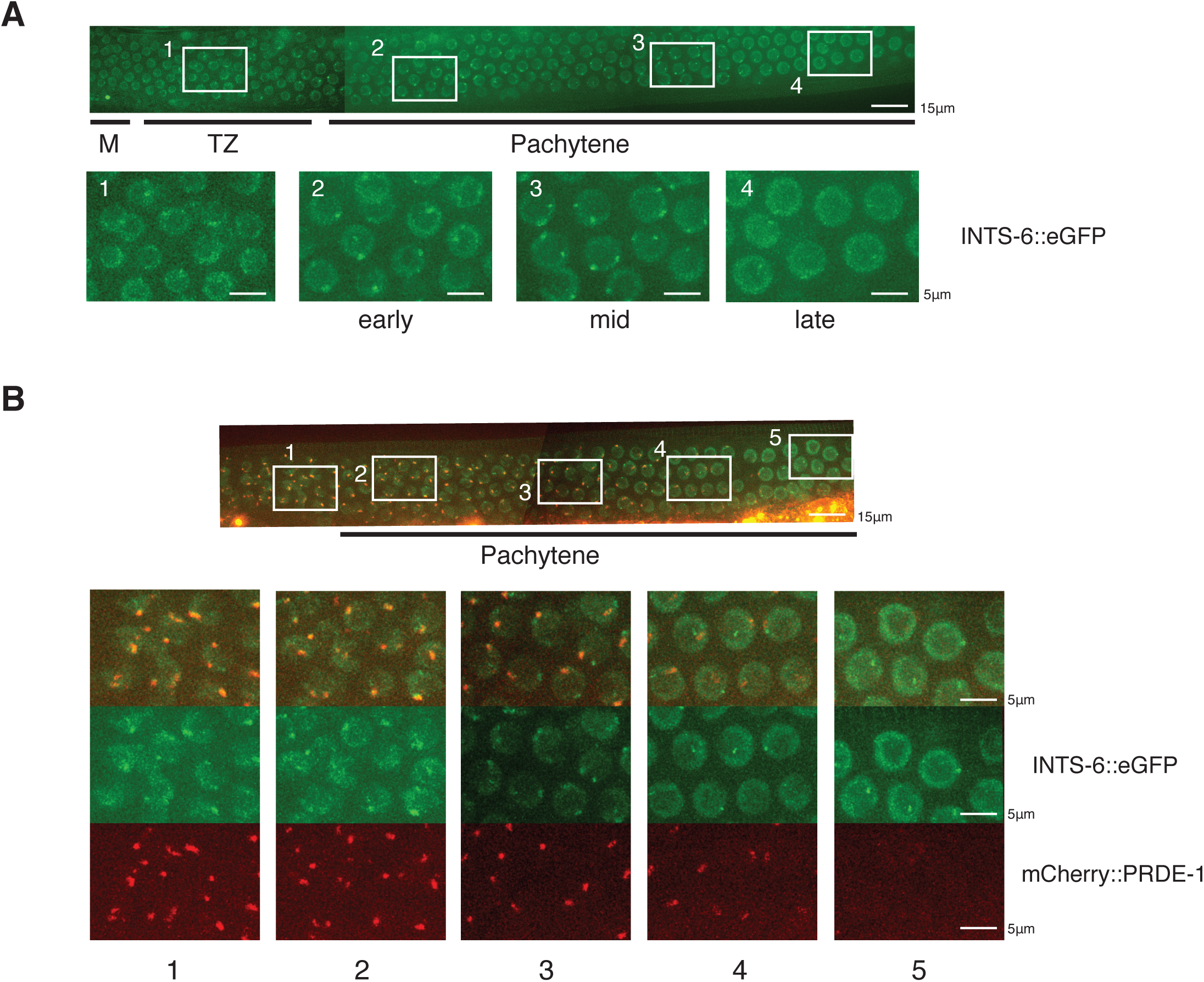
Integrator localizes to sites of piRNA biogenesis in germ cells. A. Live imaging of a rescue transgene expressing an eGFP-tagged version of Integrator subunit 6 (INTS-6) in the germline of *C. elegans*. M=mitotic region, TZ=transition zone. B. Co-localization between INTS-6::eGFP and mCherry::PRDE-1 in germ cells. PRDE-1 and INTS-6 co-localize in early stages of meiosis (1,2) and up to mid-pachytene (3), where an additional INTS-6 focus appears (3). When PRDE-1 foci are about to disappear, co-localization is lost (4), and the additional INTS-6 focus remains present well after the loss of PRDE-1 foci (5).

### Chromatin fractionation reveals two distinct populations of nascent piRNA precursors

We recently described a sequence signature of increased AT content downstream of motif-dependent 21U-RNAs resulting in a low melting temperature region (hereforth referred to as “termination signal”) (Beltran et al., 2019). We showed that the strength of this signal correlates with the length of motif-dependent piRNA precursors, and suggested that it contributes to termination by promoting pausing of early elongating Pol II due to the low stability of the AT-rich nascent RNA-DNA hybrid. We also showed that the elongation factor TFIIS, which is involved in the rescue of backtracked Pol II complexes, is required for efficient motif-dependent piRNA biogenesis (Beltran et al., 2019). On the basis of these observations, we hypothesized that Integrator might function in motif-dependent piRNA biogenesis by processing nascent RNA associated with promoter-proximal Pol II, terminating transcription.

To test this model, we fractionated purified germ nuclei into chromatin bound and nucleoplasmic fractions. This allowed us to distinguish chromatin-bound nascent piRNA precursors and nucleoplasmic precursors released from chromatin. We generated short capped RNA libraries up to an insert size of 75 bp from both fractions (Figure 3A). We observed longer precursors in the chromatin-bound fraction compared to nucleoplasm, suggesting that this approach indeed captures earlier piRNA precursors (Supp. Fig. 3A).

**Figure 3.**
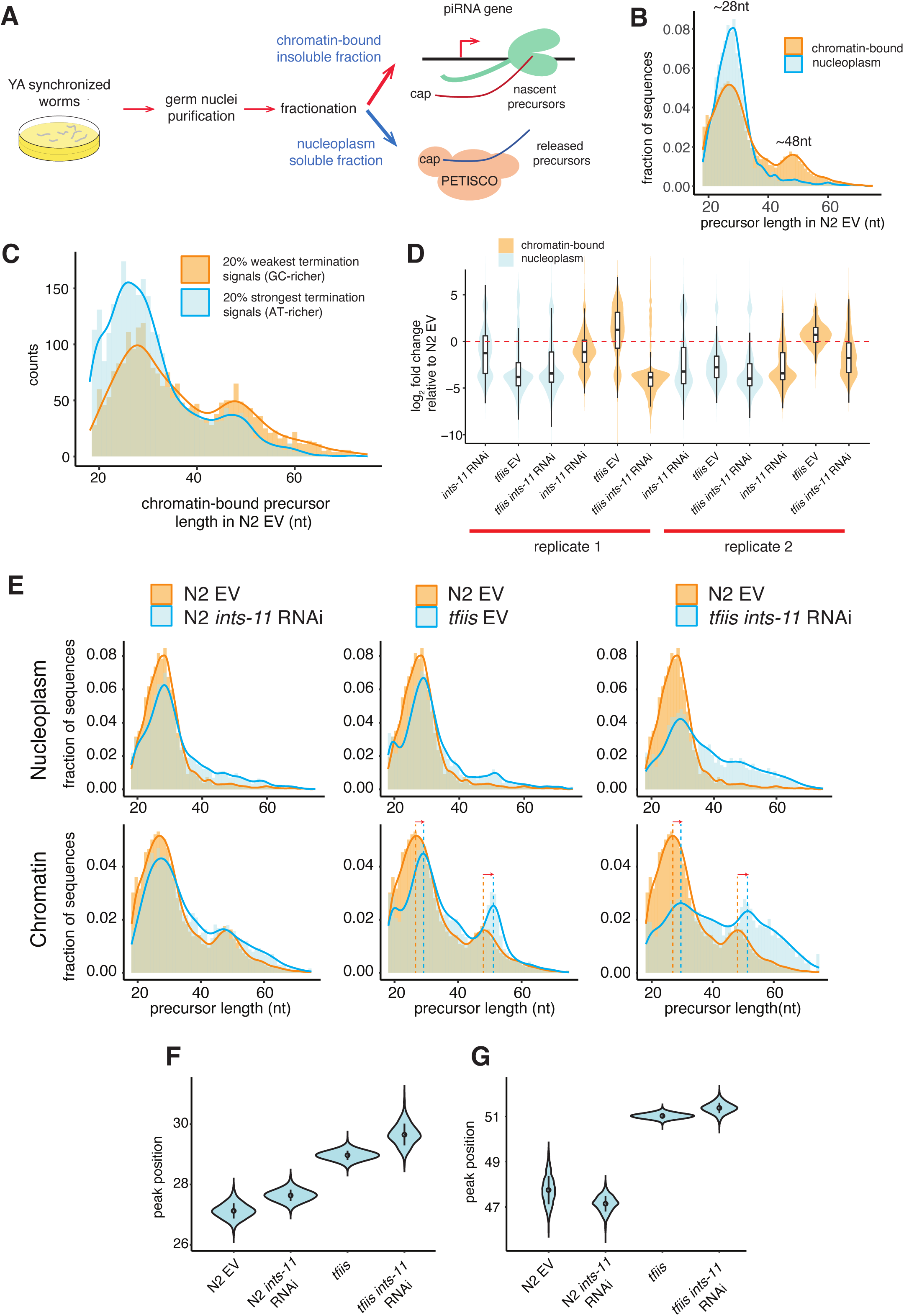
Chromatin fractionation identifies nascent piRNA precursors. A. Experimental design. Germ nuclei isolated from empty vector or *ints-11* RNAi-treated nematodes were fractionated into chromatin-bound and nucleoplasmic fractions. RNA corresponding to each fraction was purified and used as input for short capped RNA libraries. B. Length distribution of chromatin-bound and nucleoplasmic piRNA precursors in wild type N2 nematodes. C. Length distribution of piRNA precursors generated from loci with weak and strong AT content enrichment downstream of the 21U-RNA. D. Distributions of log2 fold change in piRNA precursor abundance in nucleoplasmic and chromatin fractions relative to N2 empty vector controls, for N2 *ints-*11 RNAi, *tfiis* empty vector, and *tfiis ints-*11 RNAi nematodes. Significance for each distribution to be different from no change was tested using a two-tailed paired Wilcoxon rank sum test, with all p-values<2.2e-16. E. Length distributions of nucleoplasmic and chromatin-bound piRNA precursors in N2 empty vector, N2 *ints-11* RNAi, *tfiis* empty vector, and *tfiis ints-11* RNAi nematodes. Length shifts in nascent RNA in *tfiis* mutants are indicated with dashed vertical lines and arrows. F-G. Distribution of the position of the ∼28 nt (F) and ∼48 nt (G) nascent RNA peaks in 2000 bootstrapped subsamples of 3000 precursors. Sampling probability was weighted by precursor abundance.

In wild type animals, chromatin bound nascent RNA followed a bimodal distribution, with a major peak at ∼28 nt and a less abundant peak at ∼48 nt. The ∼48 nt peak was not present in nucleoplasmic RNA (Figure 3B). Weaker termination signals (lower AT content) downstream of the 21U-RNA locus were associated with a larger proportion of sequences mapping to the 48 nt peak (Figure 3C). Upon *ints-11* knockdown, we observed a decrease in nucleoplasmic and chromatin-bound precursor levels (Figure 3D), and an increase in length in both fractions (Figure 3E). Mutation of *C. elegans* TFIIS (T24H10.1) led to a decrease in nucleoplasmic precursor levels and an increase in their length (Figure 3D,E). However, chromatin-bound precursors had increased levels in *tfiis* mutants, and showed an accumulation of nascent RNAs at the ∼48 nt peak, suggesting that TFIIS activity is important for termination of Pol II at the 48 nt peak. Knockdown of *ints-11* in *tfiis* mutants led to a further increase in precursor length in both nucleoplasmic and nascent fractions (Figure 3E). Interestingly, the position of both nascent RNA peaks was shifted in *tfiis* mutants, from 27 to 29-30 nt and from 47-48 to 51 nt. This shift persisted upon *ints-11* knockdown in a *tfiis* mutant background, indicating that it is not Integrator-dependent (Figure 3E,F,G). These differences in length are likely to result from reduced efficiency of cleavage of the 3’ end of nascent RNAs by Pol II after pausing and backtracking, suggesting that both peaks indeed correspond to Pol II pause sites on chromatin.

### Integrator cleaves nascent piRNA precursors associated with promoter-proximal Pol II

On the basis of our preceding observations, we hypothesised that Pol II pauses after it has transcribed ∼28 nt, due to increased AT content found downstream of 21U-RNAs (Beltran et al., 2019). The paused polymerase could be resolved either through release from DNA, resulting in release of a ∼28 nt piRNA precursor, or by Pol II resuming before arresting again ∼48 nt downstream of the TSS. Since ∼28 nt nascent RNAs are contained within the RNA exit channel of Pol II, termination of Pol II at the ∼28 nt peak would be independent of Integrator. Termination at the ∼48 nt peak, in contrast, would result from Integrator cleavage. This model would account for the increased proportion of ∼48 nt nascent RNAs upon Integrator knockdown, and the small but consistent 3’ shift in the position of the two peaks in the absence of TFIIS.

To test this model, we searched for cleavage products with a 5′ monophosphate that might result from Integrator cleavage of nascent RNA (Figure 4A). After removal of piRNA reads, we observed a population of 5′ monophosphate fragments with 5′ ends peaking at +38 nt of piRNA TSSs, which we designated as putative cleavage products (Figure 4B). Interestingly, *ints-*11 knockdown resulted in a clear decrease in cleavage products (Figure 4B,C, Supp. Fig. 4A,B), suggesting that these result from Integrator cleavage. These fragments had a median length of 20 nt (Figure 4D), with their 3′ ends centered 58 nt downstream of piRNA TSSs (Supp. Fig. 4C). This data suggests that Integrator processing occurs when Pol II has transcribed approximately 10 nt past the +48 pause site, resulting in the release of 38 nt capped piRNA precursors from chromatin. TFIIS mutation alone also resulted in a reduction in Integrator cleavage products (Figure 4B,C, Supp. Fig. 4A,B). This indicates that TFIIS activity is important for Integrator-mediated processing of nascent RNAs, by promoting Pol II transcription through the +48 pause site. This model is consistent with the accumulation of ∼48 nt nascent precursors in *tfiis* mutants (Figure 3D,E). Similarly, *ints-11* knockdown in a *tfiis* background resulted in a reduction in cleavage products (Figure 4B,C, Supp. Fig. 4A,B).

**Figure 4.**
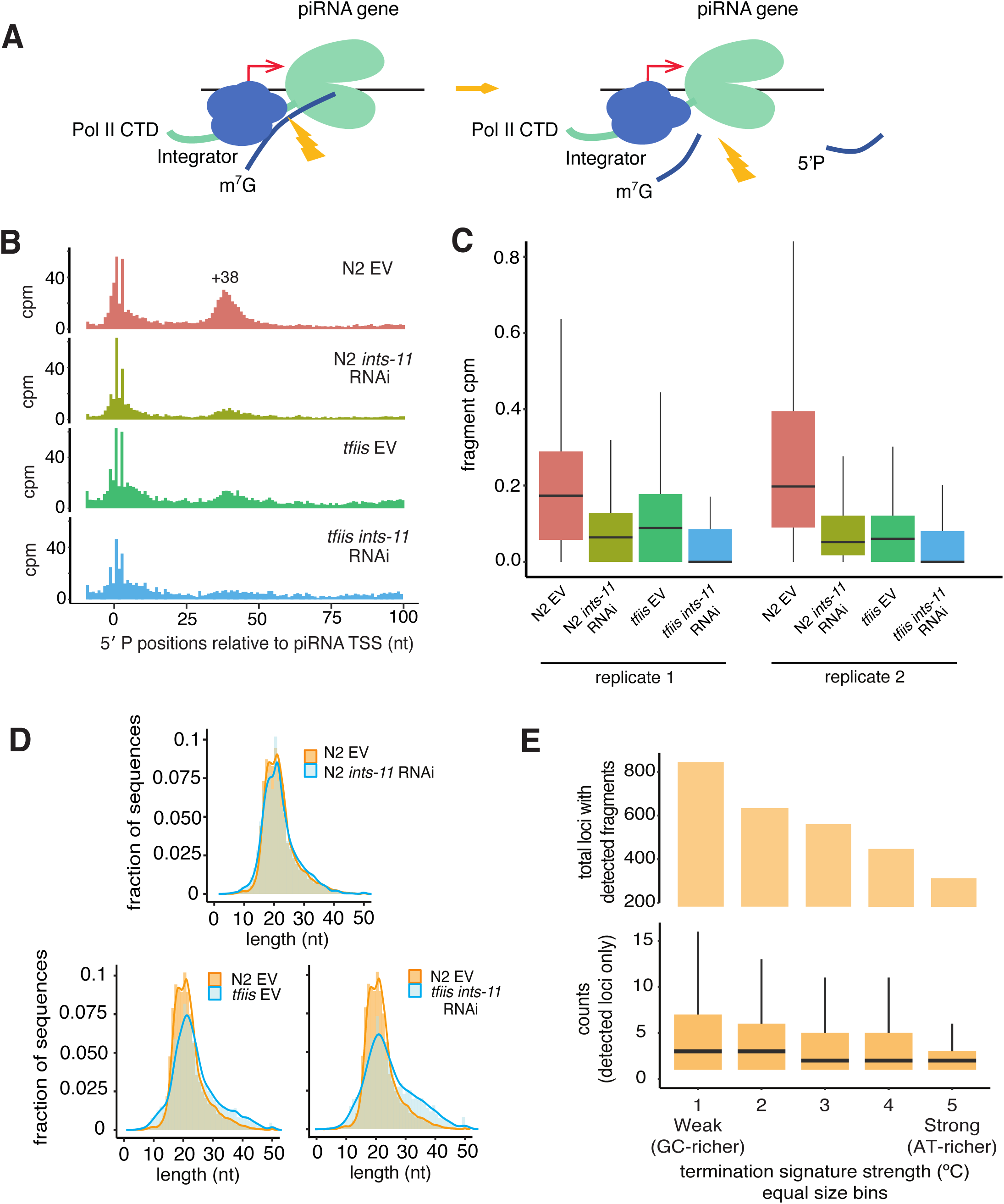
Integrator-mediated cleavage of nascent piRNA precursors generates short capped RNAs. A. Model for Integrator cleavage of nascent piRNA precursors associated with promoter-proximal Pol II. Cleavage results in the production of a short capped RNA, and a 3’ degradation fragment. B. Signal of 5′ P small RNA 5′ ends mapping to piRNA loci as a function of the distance to piRNA TSSs, after removal of reads corresponding to mature 21U-RNAs (see Methods). The signal is normalized to counts per million of non-structural mapped reads. A peak of 5′ P ends centered at +38 from piRNA TSSs is observed. C. Distributions of counts per million of 5′ P ends mapping from +25 to +50 of piRNA TSSs across the different genotypes and conditions. Two independent biological replicates are shown. D. Length distribution of putative 3′ degradation fragments with 5′ P ends mapping at +28-+58 of the 5′ U of annotated 21U-RNAs. E. Number of loci with detected cleavage fragments in N2 empty vector nematodes across percentile bins of increasing termination signal strength (top panel). Distributions of fragment read counts per locus for detected loci across percentile bins of increasing termination signal strength (bottom panel).

An important test of our model is that stronger AT-rich signals would result in reduced reliance on Integrator cleavage due to a larger proportion of termination events happening at the 28 nt peak. In support of this, loci with stronger AT-rich signals tended to have reduced abundance of cleavage products (Figure 4E, Supp. Fig. 4D). This data suggests that the AT-rich signal affects pausing and progression of Pol II to the 48 nt peak, and not Integrator processing efficiency. Consistently, the differences in nascent piRNA precursor length between loci with strong and weak termination signals were still present upon knockdown of *ints-11*, suggesting that *ints-11* does not mediate the effect of the termination signal itself (Supp. Fig. 5).

### Pol II elongation control is independent of Integrator activity at the majority of piRNA loci

Having established that Integrator cleaves nascent piRNA precursors associated with paused Pol II, we set out to test whether Integrator acts to prevent further Pol II elongation beyond the second pause site at 48 nt downstream of the piRNA TSS. To investigate this, we performed super low-input carrier CAGE (Cvetesic et al., 2018) using chromatin-bound RNA as input material and subjected the libraries to paired-end sequencing. This approach captures 5′ ends of nascent long (>200 nt) capped RNAs, allowing us to identify putative piRNA precursor initiating specifically 2 nt upstream of 21U-RNAs. Paired-end sequencing allowed us to obtain lower-bound estimates of their length from the position of the 3’ end of the fragments generated through random priming.

Nascent RNA libraries had much higher intron retention rates than a total RNA control (Supp. Fig. 6A,B), validating that chromatin fractionation captures nascent RNAs. Across samples, we observed an enrichment of unique CAGE tags initiating 2 nt upstream of motif-dependent 21U-RNAs corresponding to 659 loci in total (3% of Ruby motif-dependent piRNAs, Figure 5A, Supp. Fig 6C). These transcripts were found at low levels in N2 EV controls, as well as in *tfiis* mutants; however, *ints-11* knockdown led to an increase in their abundance (Figure 5B,C), both in an N2 background (2.83 median fold increase), and a *tfiis* background (22.62 median fold increase) (Figure 5B). Importantly, this effect was highly specific for CAGE reads initiating 2 nt upstream of piRNAs (Figure 5C), confirming that this signal captures transcription from piRNA promoters. These transcripts peaked at a length of ∼400-500 nt (Figure 5D), similarly to the library insert size, suggesting that this is a lower-bound estimate of their length. Thus *ints-*11 knockdown leads to an increase in readthrough transcription at piRNA loci.

**Figure 5.**
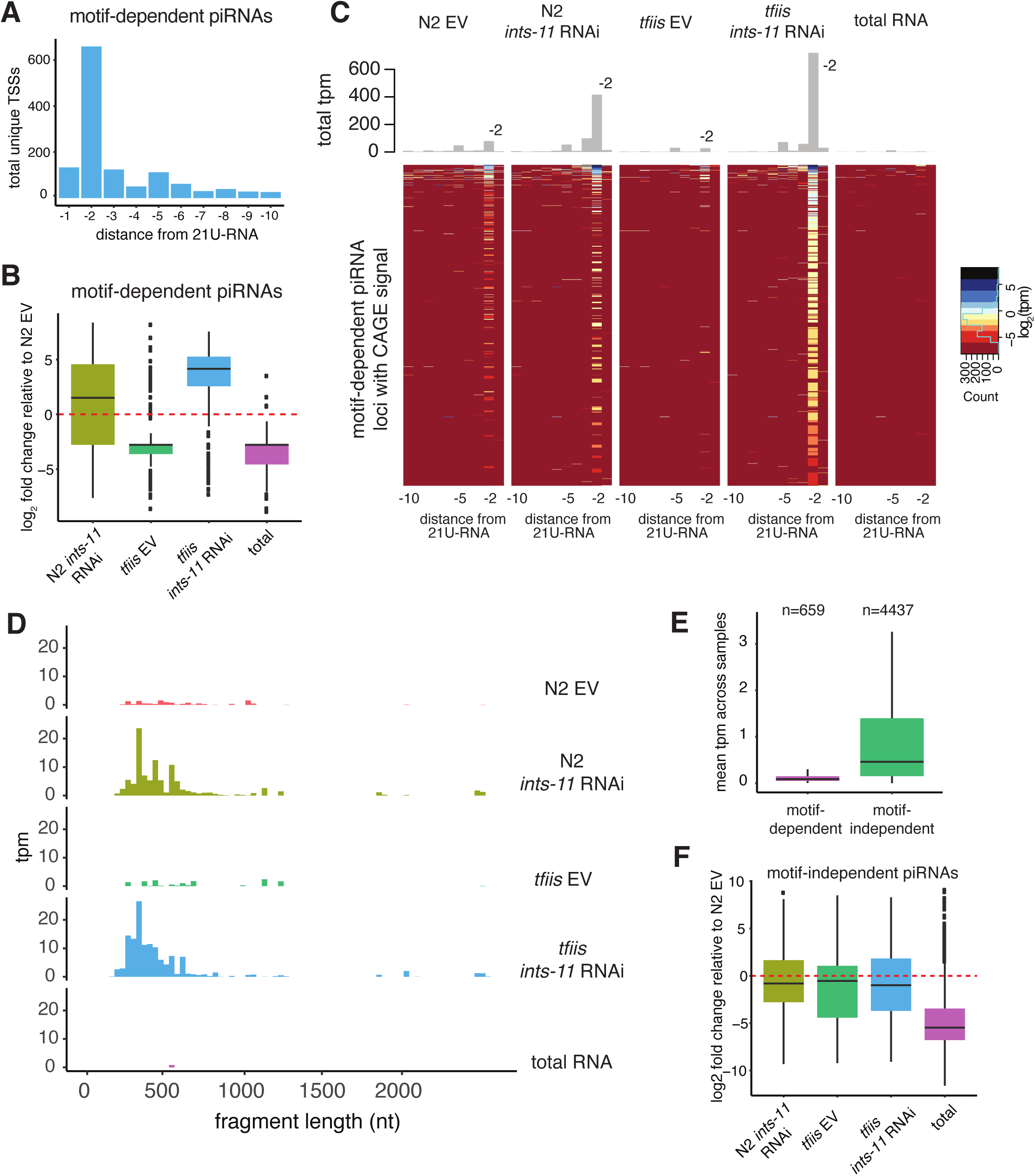
Integrator knockdown results in transcriptional readthrough in a subset of motif-dependent piRNA loci. A. Number of unique CAGE-supported TSSs mapping to a 10 nucleotide region upstream of motif-dependent 21U-RNAs. A clear enrichment of TSS 2 nt upstream of 21U-RNAs is observed. B. Distributions of log_2_ fold change in CAGE signal corresponding to 2 nt upstream motif-dependent piRNAs upon *ints-*11 RNAi knockdown in an N2 and a *tfiis* mutant background. N2 empty vector was used as a baseline for fold change calculation. C. Heatmap showing changes in CAGE signal upstream of motif-dependent 21U-RNAs upon *ints-*11 knockdown in an N2 and a *tfiis* mutant background. The total cpms corresponding to each nucleotide position upstream to 21U-RNAs are shown as a barplot (top part). D. Length distributions of CAGE fragments initiating 2 nt upstream of motif-dependent 21U-RNAs. The total counts in each length bin are normalized to counts per million of mapped reads. E. Average normalized CAGE signal (transcripts per million) 2 nt upstream of motif-dependent and motif-independent piRNAs (excluding undetected loci). F. Distributions of log_2_ fold change in CAGE signal corresponding to 2 nt upstream motif-independent piRNAs upon *ints-*11 RNAi knockdown in an N2 and a *tfiis* mutant background. N2 empty vector was used as a baseline for fold change calculation.

Long capped RNAs initiating 2 nt upstream of motif-independent piRNAs were much more abundant, with 4437 motif-independent piRNA loci producing detectable long capped RNAs (45% of motif-independent loci) (Figure 5E, Supp. Fig. 6D,E). Their abundance, however, was slightly reduced upon Integrator knockdown, in contrast to motif-dependent piRNAs (Figure 5F).

Despite the clear increase in readthrough transcription at motif-dependent piRNA loci upon *ints-11* knockdown relative to WT, we still only detected such events at a minority of loci (4.65%). This observation is not likely to result from low sensitivity of CAGE since the levels of short piRNA precursors are not significantly different between CAGE-detected and undetected piRNAs (Supp. Fig. 6F); and the number of detected motif-dependent piRNAs is saturated at the sequencing depth we obtained for our CAGE libraries (Supp. Fig. 6G). In order to better understand the factors influencing Pol II promoter escape at piRNA loci, we examined chromatin accessibility (Jänes et al., 2018) and H3K27ac levels (Han et al., 2019) at piRNA promoters. Motif-dependent piRNA promoters showed extremely low accessibility levels below genome-wide average (Supp. Fig. 7A); however, a clear increase in ATAC signal is observed around piRNA TSSs when comparing wild-type adult worms to *glp-1* mutants lacking a germline (1.5-fold mean fold increase) (Supp. Fig. 7B). Motif-independent piRNAs, in contrast, showed on average 10 times higher accessibility than genome-wide average, both in wild type and *glp-1* nematodes (Supp. Fig. 7A,B). Both types of piRNA loci also showed clear differences in the levels of H3K27ac: motif-dependent loci showed extremely low levels, while motif-independent loci were highly enriched; this enrichment was especially high in the germline (Supp. Fig. 7C). The subset of motif-dependent piRNA promoters that were detected by CAGE tended to have slightly higher levels of H3K27ac and accessibility, but these are still very low relative to motif-independent loci (Supp. Fig. 7D,E). We thus conclude that the differences in readthrough transcription between motif-dependent and motif-independent piRNA promoters correlate with chromatin accessibility and H3K27ac status. However, within motif-dependent loci, accessibility and H3K27ac levels do not explain differences in readthrough.

This scenario contrasts to snRNA loci, where *ints-*11 knockdown led to a dramatic increase in CAGE signal at all loci (Supp. Fig. 8A,B), reflecting extensive readthrough past snRNA termination sites, which has been described to result in fusion transcripts with downstream protein-coding genes (Figure 6, Gómez-Orte et al., 2019).

**Figure 6.**
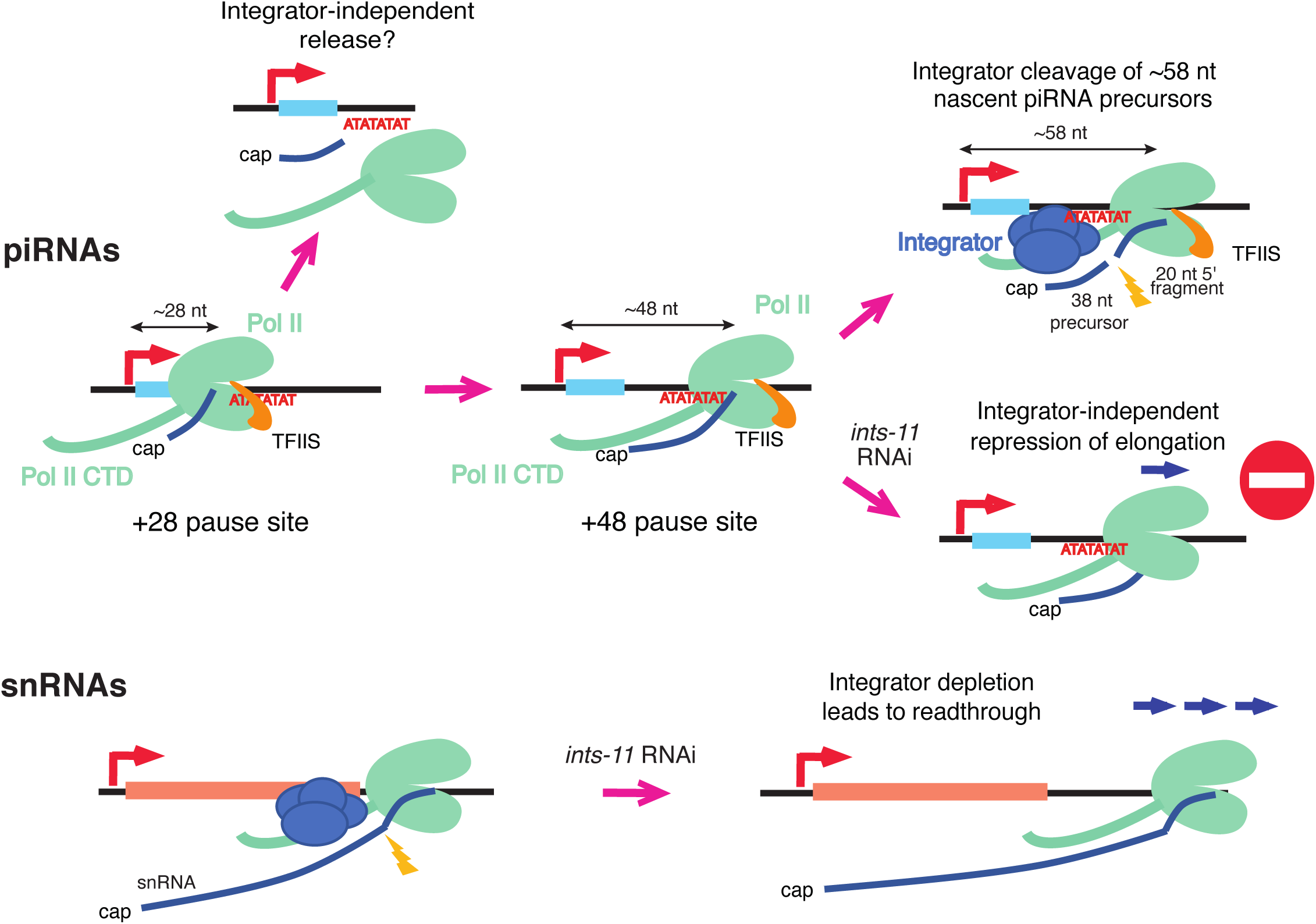
Model for the role of Integrator in piRNA transcription termination. At snRNA loci, Integrator generates snRNA 3′ ends leading to Pol II termination. *ints-*11 knockdown results in readthrough past termination sites and polyadenylation of snRNA transcripts. At piRNA loci, Pol II pauses ∼28 nt and ∼48 nt downstream of piRNA TSSs. ∼28 nt short capped RNAs may be generated via direct dissociation of Pol II from chromatin at the ∼28 nt pause site. When Pol II transcribes past the ∼48 nt pause site, Integrator acts to cleave nascent precursors approximately when Pol II is at +58 from piRNA TSSs, generating ∼38 nt short capped RNAs and ∼20 nt degradation fragments. *ints-*11 knockdown does not lead to Pol II readthrough, suggesting that additional mechanisms prevent Pol II elongation at piRNA loci.

### Integrator attenuates protein-coding gene expression in C. elegans

We assessed the role of Integrator in protein-coding gene transcription by examining the changes in short capped RNA and CAGE reads initiating at gene promoters (Chen et al., 2013). We identified a total of 437 genes with changes in short capped RNA signal (FDR<0.1; 371 upregulated, 66 downregulated, Supp. Fig. 8C), as well as 470 genes with changes in CAGE signal (FDR<0.1; 330 upregulated, 140 downregulated, Supp. Fig 8D). In addition, changes in short and long capped RNAs tended to correlate well, with very few genes showing changes in the opposite direction (Supp. Fig. 8E-G). The enrichment of upregulated genes suggests that Integrator-mediated termination negatively regulates transcription of a subset of protein-coding genes, as has been recently reported (Elrod et al., 2019). Since motif-independent piRNA loci tend to localize within protein-coding gene promoters genome-wide, we wondered whether they may be generated as a by-product of premature Pol II termination at protein-coding genes by Integrator. Motif-independent piRNA loci, however, were not enriched in the promoters of genes that become upregulated upon Integrator knockdown (Supp. Fig. 8H), consistent with an Integrator-independent biogenesis pathway for motif-independent piRNAs.

## Discussion

In this work, we gain insight on the fundamental mechanism of piRNA biogenesis in *C. elegans*, and explore the role of the Integrator complex in the termination of promoter-proximal Pol II at piRNA loci.

### Parallels between snRNA and piRNA transcription

Multiple lines of evidence suggest that piRNA promoters evolved from snRNA promoters (Beltran et. al., 2019, Domingues et. al., 2019, Weng et. al., 2019). Here, we explore the parallels between the mechanisms of piRNA and snRNA transcription. We show that the Integrator complex is recruited to sites of piRNA biogenesis upon piRNA transcription activation. It is unclear whether Integrator recruitment to piRNA promoters is directly mediated by the USTC complex (SNAPc/PRDE-1/TOFU-5), or whether Integrator is recruited to the initiating Pol II independently of SNAPc. Integrator has been shown to associate with the CTD of Pol II phosphorylated on Ser7 (Egloff et al., 2007, 2010). It will be of interest to explore whether Ser7-P is associated with piRNA biogenesis, and the mechanisms directing this modification. The presence of piRNA-independent Integrator foci may reflect Integrator-dependent regulation of snRNA genes. Indeed, snRNA genes have been shown to spatially interact and coalesce into Cajal bodies (Frey and Matera, 2001; Wang et al., 2016). This phenomenon may share some mechanistic aspects with the formation of piRNA foci in the *C. elegans* germline.

### Two distinct populations of piRNA precursors

By isolating chromatin-bound RNA from germ nuclei, we characterize two distinct populations of nascent piRNA precursors; one peaking at ∼28 nt, and a second population of ∼48 nt long, representing two Pol II pause sites on chromatin. In addition, we provide evidence that Integrator acts to cleave 53-63 nt nascent precursors past the +48 pause site, resulting in the release of 33-43 nt short capped piRNA precursor transcripts (Figure 6). Several studies had set out to address the length of piRNA precursors in the past. Cecere et al., 2012 reported the existence of a ∼75 nt long precursor detected by RACE, but high-throughput sequencing-based approaches detected 26-28 nt precursors (Gu et al., 2012; Weick et al., 2014). Our discovery of a second population of 40-60 nt nascent precursors reconciles these previous observations by suggesting that longer precursor transcripts, whilst rare, can be detected at piRNA loci.

Interestingly, a large fraction of nascent precursors peak at a length of ∼28 nt, as observed for nucleoplasmic precursors. This raises the question of whether these are truly chromatin-bound. We previously suggested that increased AT content downstream of 21U-RNAs leads to Pol II pausing ∼28 nt downstream of the TSSs (Beltran et al., 2019). Consistent with this model, TFIIS mutation results in a 2-3 nt shift in both +28 and +48 peaks, suggesting pausing and backtracking of Pol II at both sites (Figure 6). In addition, Integrator cleavage fragments are reduced in loci with high AT content downstream of 21U-RNAs, suggesting that Pol II is less likely to reach the +48 peak in these loci. Pol II may be released from the +28 pause site independently of Integrator, and Integrator may act as a failsafe termination mechanism when transcription continues beyond +48 (Figure 6). A recent study also supports the existence of two Pol II pause sites at protein-coding genes in human cell lines (Aoi et al., 2020). Interestingly, accumulation of Pol II at the second pause site is observed upon NELF depletion (Aoi et al., 2020). *C. elegans* does not have NELF (Maxwell et al., 2014) potentially favouring the transition of Pol II to the +48 peak.

### Integrator and Pol II elongation control

Our analysis of the role for Integrator at piRNA loci in *C. elegans* fits with a recent study demonstrating that Integrator terminates promoter-proximal Pol II at protein-coding gene promoters to attenuate gene expression (Elrod et al., 2019). The short-capped RNAs produced as a by-product of Integrator cleavage at protein-coding gene promoters are rapidly degraded by the nuclear exosome. At *C. elegans* piRNA loci, instead, the resulting short capped RNAs are handed in to a specialized RNA processing complex resulting in mature piRNA production (Cordeiro Rodrigues et al., 2019; Zeng et al., 2019). How piRNA precursors are recognized and channelled into the downstream processing pathways while avoiding nuclear RNA surveillance will be interesting to explore in the future.

Our data reveals an interesting interplay between TFIIS and Integrator function in the regulation of promoter-proximal Pol II. TFIIS mutation results in increased nascent precursors, and an accumulation of ∼48 nt precursors, but a decrease in degradation fragments, suggesting that TFIIS activity modulates the ability of Integrator to process nascent RNAs. The observation that Integrator processing occurs downstream of the +48 pause site suggests that the role of TFIIS is to promote elongation through the +48 site prior to Integrator termination. Interestingly, *C. elegans* mutants of the RPB-9 subunit of Pol II exhibit defects in piRNA biogenesis similar to *tfiis* mutants (Berkyurek et al., cosubmitted). This data is consistent with previous studies indicating that TFIIS binding to Pol II is compromised in Δ*rpb-9* mutants in *Saccharomyces cerevisiae* (Sigurdsson et al., 2010). Indeed, the capacity of TFIIS to promote Pol II transcription through arrest sites *in vitro* depends on RPB-9 (Awrey et al., 1997). Understanding how the regulation of Pol II early elongation influences the ability of Integrator to prematurely terminate transcription will be an interesting avenue for future study.

At snRNA loci, Integrator is required to terminate transcription and in its absence substantial readthrough transcription is observed (Baillat et al., 2005; Gómez-Orte et al., 2019). In our study, we observed similar evidence of readthrough transcription after Integrator knockdown at snRNA loci. In contrast, Integrator depletion did not result in readthrough transcription at most motif-dependent piRNA loci. This indicates that the cleavage activity of Integrator is not required to prevent elongation at motif-dependent piRNAs. Pol II elongation control is thus likely to operate independently of Integrator via additional mechanisms (Figure 6). One possibility is that elongation repression is chromatin-based. Motif-dependent piRNAs localize to broad domains of H3K27me3 (Beltran et al., 2019), the vast majority of which also have low accessibility and low H3K27 acetylation (Supp. Fig. 8A-D), reminiscent of poised enhancers (Creyghton et al., 2010). In contrast, motif-independent piRNAs, which show extensive readthrough, are in highly accessible H3K27ac-rich genomic regions. The tight control of Pol II elongation at motif-dependent piRNA loci may be a consequence of evolutionary pressure to avoid transcriptional interference between neighbouring piRNA loci given the high density of loci in piRNA cluster regions that are simultaneously activated.

### Integrator: piRNA functions beyond nematodes?

The involvement of Integrator in piRNA biogenesis adds to the existing evidence that snRNA and nematode piRNA transcription are functionally and evolutionarily related. Whether this principle extends to piRNAs in other organisms remains unknown; however, the piRNA biogenesis factor cutoff has been shown to bind to snRNA promoters in *Drosophila melanogaster* (Pritykin et al., 2017). Interestingly, Integrator was identified in genetic screens to be required for piRNA-mediated silencing of a germline reporter in *Drosophila* (Handler et al., 2013), making Integrator a possible candidate to assist in 3’ end formation of piRNAs in *Drosophila* and mouse. Our data adds to the growing evidence indicating co-option of functional modules from a range of existing transcriptional and RNA metabolic processes in metazoan piRNA biogenesis, and expands the repertoire of non-coding RNAs under Integrator control in metazoans.

## Methods

### Nematode culture and RNAi

*C. elegans* nematodes were grown at 20°C in standard nematode growth medium (NGM) agar plates feeding on OP50 *E. coli*. RNAi clones from the Ahringer library (Kamath and Ahringer, 2003) were grown overnight in LB supplemented with 50 μg/ml ampicillin and 25 μg/ml tetracycline. The next morning, cultures were diluted in LB 50 μg/ml ampicillin, grown to an OD of 0.5-0.6, and seeded into 1 week old NGM plates supplemented with 1 mM IPTG and 50 μg/ml ampicillin. Plates were dried for 48 h before plating synchronized L1 worms obtained by hypochlorite treatment followed by 24 h starvation in M9 media.

### Live imaging

Live imaging of SX1316 piRNA sensor worms, as well as INTS-6::eGFP::3xFLAG and PRDE-1::mCherry transgenes was carried out in a DeltaVision fluorescence microscope system. Worms were picked into a 2 μl drop of M9 with 0.5 μM levamisole on a slide, and subsequently immobilized in a hydrogel matrix with microbeads as described in Dong et al., 2018. Live imaging was carried out within 15 min of slide preparation.

### Immunofluorescent staining

Mouse anti-FLAG M2 monoclonal antibody (F1804, Sigma) preadsorbed with *C. elegans* lysates to remove background was used for immunofluorescent staining. 15-20 worms were picked onto 15 μl M9 Tween 0.1% on a poly-lysine-coated slide, and dissected using a gauge needle. Another 15 μl 2% paraformaldehyde in M9 Tween 0.1% were added, some solution was removed and a coverslip was placed on top to let the germlines fix for 5 minutes. The slides were then frozen in liquid nitrogen, worms were cracked by quickly removing the coverslip, and slides were fixed in - 20°C methanol for at least 1 minute. Slides were then washed by immersion in PBST for 5 min, 3 times, and blocked with PBST with 1% BSA for 1 h at room temperature. Slides were then incubated with mouse anti-FLAG M2 antibody diluted 1:500, and rabbit anti-mCherry antibody (GTX128508) diluted 1:500 in blocking solution, in a wet chamber, overnight, at 4°C. The next day, slides were washed in PBST for 10 minutes, 3 times, and then incubated with goat anti-mouse Alexa 488 secondary antibody at 1:200 dilution, and goat anti-rabbit Alexa 594 secondary antibody at 1:200 dilution in PBST for 1-2 h in the dark. Slides were again washed in PBST for 10 minutes, 3 times, and incubated with 100 μl DAPI 1 μg/ml for 5 minutes in the dark. Slides were finally washed in PBST in the dark for 20 minutes. Excess liquid was removed and slides were mounted with Vectashield and sealed with nail polish.

### RNA extraction

For total RNA whole animal samples, young adult synchronized populations of worms were grown and washed with M9 three times to remove bacterial residues. For each 100 μl of worm pellet, 1ml of Trizol was added. 5 cycles of freeze-cracking were carried out, freezing in liquid nitrogen, followed by thawing in a water bath at 37°C. Tubes were vortexed for 30 s on, 30 s off, for 5 minutes, and incubated for 5min at room temperature. 200 μl chloroform per ml of Trizol were added, tubes were shaked vigorously, incubated 2-3 min at room temperature, and centrifuged at full speed for 10 min at 4°C. The top layer was transferred to a new tube, and RNA was precipitated overnight at −20 °C with 1 μl glycogen and an equal volume of isopropanol.

### Germ nuclei purification and chromatin fractionation

Germ cells were purified similarly to the protocol described in Han et al., 2019. Approximately 50 100 mm standard NGM or RNAi plates of synchronized young adult worms were used per sample. Worms were washed off plates with M9 and collected in a 15 ml conical tube, and washed several times with M9 to clean from bacteria. The final wash was done with ice cold nuclear purification buffer (NPB: 10 mM HEPES pH 7.6, 10 mM KCl, 1.5 mM MgCl_2_, 1 mM EGTA, 0.25 M sucrose, 0.025% TritonX-100, 50 mM NaF, 1 mM DTT, 40 mM β-glycerol, 2 mM Na_3_VO_4_). Worms were resuspended in NPB and transferred to a previously chilled metal grinder (#08-414-20A), and 1-3 strokes were applied, monitoring the extent of homogenization after each stroke, such that no more than 10-15% of worms were broken. The solution was transferred to a prechilled 50 ml conical tube, and vortexed for 30 s at medium speed, then incubated on ice for 5 min, twice. The sample was then filtered through a 100 μm nylon mesh filter (Falcon), followed by three 40 μm (Falcon) and three 20 μm filters (PluriSelect). The filtered solution was recovered and centrifuged at 100 g for 6 min at 4°C to pellet debris (usually a good germline yield has little or no material pelleted in this step). The supernatant was recovered and centrifuged at 2500 g for 10 min at 4°C to pellet nuclei, and the supernatant was discarded. Isolated nuclei were then subjected to chromatin fractionation as described in Nojima et al., 2016. Briefly, nuclei were resupended in 67.5 μl of ice-cold NUN1 buffer (20 mM Tris-HCl pH=7.9, 75 mM NaCl, 0.5 mM EDTA, 50% glycerol v/v, 1x cOmplete protease inhibitor cocktail), and mixed gently by pipetting. 0.6 ml of ice-cold NUN2 buffer (20 mM HEPES pH=7.6, 300 mM NaCl, 0.2 mM EDTA, 7.5 mM MgCl_2_, 1% v/v NP-40, 1 M urea, 1x cOmplete protease inhibitor cocktail) were added. The mix was vortexed at maximum speed and incubated on ice for 15 min, vortexing every 3-4 during the incubation to precipitate the chromatin fraction. The mix was then centrifuged at 16000 g for 10 min at 4°C. The supernatant (nucleoplasm fraction) was collected, split into two aliquots of 300 μl, and 900 μl Trizol LS were added to each aliquot and mixed by vortexing to proceed with RNA extraction. To isolate chromatin-bound RNA, the chromatin pellet was resuspended in 200 μl HSB buffer (10 mM Tris-HCl pH=7.5, 500 mM NaCl, 10 mM MgCl_2_) with 10 μl TURBO DNase (stock 2 U/μl) and 10 μl Superase-In RNase inhibitor (stock 20 U/μl), and incubated at 37°C for 10 min, vortexing to help resuspension and solubilization. 2 μl of proteinase K (stock 20 mg/ml) were then added, and the mix was incubated at 55°C for 10 min. 600 μl Trizol LS were added to proceed with RNA extraction.

### CIP-Rpph treatment of RNA and small RNA library preparation

To enrich for short-capped RNAs, we first pretreated 2 μg total RNA, or 500 ng of chromatin-bound or nucleoplasmic RNA with 2 µl Quick CIP (NEB) in a total volume of 20 µl for 90 min at 37°C. RNA was phenol-chloroform extracted, and precipitated at −20°C overnight with 1/10 vol 3 M AcNa, and 3 vol 100% EtOH and 1 µl glycogen. The next day, RNA was resuspended and treated with 7.5 units (1.5 µl) RNA 5′ pyrophosphohydrolase (NEB) for 1 h at 37°C in a total volume of 20 µl. RNA was extracted and precipitated overnight as described for CIP treatment. The resulting RNA was used as input for small RNA library preparation using the TruSeq small RNA kit (Illumina) according to the manufacturer’s instructions, except for an increase in the number of PCR cycles from 11 to 15. An insert size range of 20-70 nt was gel purified, DNA was eluted from the gel in 0.3 M NaCl, and EtOH precipitated overnight. For direct 5′ P libraries, 1 μg total RNA was used as input, and an insert size range of 20-35 nt was gel purified. Libraries were quantitated using Qubit and Tapestation prior to pooling in groups of 6 to 12 per lane, and sequenced on an Illumina HiSeq2000 (50bp single end libraries), or an Illumina NextSeq instrument (75 bp single end libraries).

### Small RNA sequencing data processing, mapping and normalization

The Illumina universal adapter was trimmed from small RNA reads using cutadapt v1.10 and reads >17nt were mapped to WS252 *C. elegans* genome (ce11) using bowtie v0.12 (Langmead et al., 2009) with parameters –v 0 –m 1. Sam files were converted to bam and bed using samtools v1.2 (Li et al., 2009) and bedtools v2.25.0 (Quinlan and Hall, 2010).

piRNA and miRNA counts were derived from direct 5′ P libraries by searching collapsed fasta files for exact matches to existing piRNA (Batista et al., 2008; Gu et al., 2012) and miRNA (Kozomara and Griffiths-Jones, 2014) annotations. Direct library sizes were estimated, using total non-structural read counts (1), using total miRNA counts (2), and extracting DEseq2 size factors from a miRNA counts table (3), resulting in similar trends. Log2 fold change in abundance was calculated for loci with more than 5 DEseq2 normalized read counts on average. 1 pseudocount was added to avoid zero counts. Additionally, the change in total mature piRNA counts relative to N2 was estimated using the same size factors.

piRNA precursor reads were identified from CIP-Rpph libraries by identifying reads mapping exactly 2 nt upstream of annotated piRNAs. Library sizes were estimated as the total counts of precursor reads mapping to previously annotated TSSs (Chen et al., 2013). The log2 fold change in abundance was calculated for loci with more than 5 reads per million of TSS-mapping scRNA reads on average. 1 pseudocount was added to avoid zero counts. Additionally, the change in total piRNA precursor counts relative to N2 was estimated using the same size factors. To estimate the effects of the termination signal strength on piRNA precursors, we calculated termination signature strength as described in Beltran et al., 2019 considering the entire transcriptional unit from the TSS to +50, and considering the region downstream of 21U-RNAs alone (+25 to +50). We plotted the length distribution of precursors across equal-sized bins of piRNA loci stratified according to pausing signature strength. We additionally plotted the length distributions for the top vs bottom 20% loci.

### Bootstrap analysis of piRNA precursor peak locations

In order to obtain robust estimates of precursor distributions, and sufficient sampling of precursors, we combined precursors identified in the two replicates for each condition. For each condition, we generated 2000 random subsamples of 3000 precursor sequences, sampling without replacement and weighting the probability of sampling by the number of reads for each precursor sequence. We fit a smoothing density curve to the distribution, identified the two local maxima of the fit corresponding to first and second peaks, and extracted the positions of the two. The resulting distributions of positions of both peaks for each condition are shown on Figure 4F,G.

### Degradation fragment analysis

Degradation fragment analysis was carried out from direct 5′ P small RNA libraries generated from RNA purified from germ nuclei, with the data processed as described above but without applying a minimum read length cutoff. Reads >15 nt initiating exactly at annotated 21U-RNA sites were discarded from the libraries, and the read coverage of the remaining 5′ P unique sequences around piRNA promoters was plotted at single nucleotide resolution in the form of an average profile and a heatmap. Additionally, the counts per million and sequences per million of 5′ and 3′ ends of 5′ P reads was similarly plotted. We found an enrichment of 21-U RNA sequences initiating within a +-5 nt window from 21U-RNA sites. These likely represent unannotated 21U-RNAs originating from overlapping TSSs from the same piRNA locus (Billi et al., 2013). We additionally removed these 21U-RNA sequences, as well as reads >15 nt initiating at those unannotated 21U-RNA sites. The remaining first nucleotide and length distributions showed an enrichment of unique sequences initiating at the −2 position from annotated 21U-RNAs. Fragments whose 5′ ends mapped in between +25 and +50 from the 5′ U of annotated 21U-RNAs, corresponding to the second peak centered at +50 were collected, and their length distribution was plotted. Total fragment counts per locus were calculated and normalized to cpm of total non-structural mapped reads. The total number of loci with detectable fragments, and well as the distribution of fragment cpms were quantified across piRNA loci binned according to the strength of their termination signals.

### SLIC-CAGE library preparation and sequencing

CAGE library preparation from chromatin-bound RNA was performed using the SLiC-CAGE protocol (Cvetesic et al., 2018, 2019). The prepared libraries were pooled and sequenced on an Illumina HiSeq2000 (100bp paired-end libraries). In order to assess the enrichment of potential piRNA precursor reads in chromatin-bound samples, a total RNA sample control was used. Intron retention ratios were estimated using IRFinder (Middleton et al., 2017), which computes a ratio between the abundance of spliced and unspliced introns for all annotated RNAs,confirming the enrichment of unspliced transcripts in nascent RNA samples.

### Analysis of SLIC-CAGE data

CAGE tags of read 1 were mapped to a reference C.elegans genome (ce11) using STAR (Dobin et al., 2013). Uniquely mapped reads were imported into R (http://www.R-project.org/) as bam files using the standard workflow within the CAGEr package (Haberle et al., 2015). Due to the template free activity of the reverse transcriptase enzyme, often a G is added at the 5’ end of the read. G’s that do not map to the genome at the 5’ end of reads are removed by CAGEr’s standard workflow. G’s that map to the genome are removed at a rate comparable to the mismatching G’s removed. The 5’ ends of reads represent CAGE-supported transcription start sites (CTSSs) and the number of tags for each CTSS reflects expression levels. Raw tags were normalised using a referent power-law distribution and expressed as normalized tags per million (TPMs) (Balwierz et al., 2009). Biological replicates were highly correlated (r^2^ > 0.9) and were therefore merged, and the distributions of log_2_ fold change in CAGE signal at the −2 position of piRNA loci were calculated. For each pair of samples, piRNA loci with non-zero counts in at least one of the conditions were included, and 0.01 tpm were added to all data points to prevent division by zero. Log_2_ transformed data for all positions upstream of piRNAs in all conditions was plotted as a heatmap, with the total sum of normalized CAGE tpms by position shown above.

For fragment length quantification, reads were aligned using using Tophat v2.1.0. (Langmead and Salzberg, 2012) with parameters -X 20000. Alignment files were sorted by read name and converted to bed12 format using pairedBamToBed12. Read pairs initiating at −2 from 21U-RNAs were retrieved and their overall length and abundance normalized to counts per million of mapped fragments was plotted. In addition, library sizes obtained from the total numbers of pair1 reads containing SL1 and SL2 spliced leader sequences were used for normalization, resulting in very similar trends to previous normalization strategies. Tophat alignments were converted to bigwig tracks using deeptools v3.1.2 bamCoverage (Ramírez et al., 2016) with parameters -bs 10 -e, tracks from the two replicates for each condition were averaged, and signal average profiles and heatmaps were generated for snRNA and protein-coding gene TSSs.

CAGE counts were randomly downsampled to multiple total count sizes (from 50,000 to 5,000,000 to match library size across libraries), and to multiple percentages of total library size (to examine saturation of piRNA detection with sequencing depth). 15 samples were obtained for each size and library, and the total number of piRNA loci with detected signal at −2 nt from the 21U-RNA were recovered. For each of the 15 subsamples, the two replicates of each library were averaged. The resulting distributions of total detected piRNAs per condition were plotted as boxplots against the depth of the subsamples.

### ATAC-seq and ChIP-seq data processing and analysis

Processed bigwig ATAC-seq tracks from young adult wild-type nematodes, and day 1 *glp-1* mutant adults were obtained from Janes et al., 2018 (GSE114439), and lifted over to ce11. Raw H3K27 acetylation ChIP-seq data from isolated germ nuclei and somatic nuclei was obtained from Han et al., 2019 (SRR9214969 to SRR9214976), mapped to ce11 using bowtie2 (Langmead and Salzberg, 2012) with default parameters, and the resulting bam files were converted to bigwig tracks using deeptools bamCoverage (Ramírez et al., 2016) with parameters -bs 1 -e 150 -- normalizeUsing RPGC --effectiveGenomeSize 96945445. Replicates were averaged using bigwigCompare, and heatmaps of H3K27ac and ATAC signal around piRNA loci were generated using deeptools computeMatrix reference-point. The signal data underlying the heatmaps was extracted in order to compare the distributions of ATAC (±100 bp around piRNA TSSs) and H3K27ac signal between groups of loci (±1 kb around piRNA TSSs).

### Differential expression analysis of gene promoters genome-wide using scRNA and CAGE count data

scRNA or CAGE TSS annotations were intersected with a set of WormBase protein-coding gene promoters (Chen et al., 2013) lifted over to ce11, in order to generate count matrices of scRNA or CAGE counts for all gene promoters genome-wide. Differential expression analysis comparing N2 empty vector chromatin bound samples to N2 *ints-11* RNAi chromatin bound samples was carried out using DESeq2. Promoters were considered to be differentially expressed using a Benjamini-Hochberg FDR threshold of 0.1. The fold changes of promoters identified as differentially expressed in either of the two experiments were extracted, and the fold change in scRNA signal was plotted against the fold change in CAGE signal. To test for enrichment of motif-independent piRNAs in each of the promoter classes, motif-independent piRNAs were first overlapped with promoters detected in the scRNA and CAGE experiments in order to determine their background enrichment. Enrichment was calculated in each promoter set as an odds ratio relative to its corresponding background; for instance the fraction of CAGE “up” promoters overlapping with motif-independent piRNAs was compared to the fraction of all CAGE detectable promoters overlapping with motif-independent piRNAs. For intersecting promoter sets, genes detected as significant in either experiment, and having a fold change greater than 2 in both experiments were selected. In these cases, background was calculated as the fraction of promoters detected in both experiments overlapping with motif-independent piRNAs.

## Acknowledgements

We would like to thank Juan Cabello for kindly sharing JCP383 ints-6::GFP nematodes with us at the beginning of the study.

## Supplementary Figure legends

**Supplementary Figure 1. Quantification of total mature piRNA and piRNA precursor levels after RNAi knockdown of *ints-*11 (relates to Figure 1)**.

A-B. Total piRNA read counts normalized using DEseq2 size factors derived from miRNA counts (A) or normalized to the total number of non-structural mapped reads (B). Normalized abundance for each pair of EV and *ints-*11 RNAi samples is shown relative to EV.

C. Average mature piRNA abundance relative to EV in EV and *ints-*11 RNAi samples normalized using miRNA-derived size factors. Error bars represent the standard error of the mean.

D. Representative examples of a silenced and a desilenced piRNA sensor transgene in the *C. elegans* germline. The proportion of animals desilencing the piRNA sensor upon empty vector and *ints-*11 RNAi treatment is shown in the right panel.

F. Total piRNA precursor counts normalized to the total number of short capped RNA reads mapping to WormBase TSSs (Chen et al., 2013). Normalized abundance for each pair of EV and *ints-*11 RNAi samples is shown relative to EV.

G. Average normalized piRNA precursor abundance relative to EV, in EV and *ints-*11 RNAi samples. Error bars represent the standard error of the mean.

H. Distributions of piRNA precursor length in *ints-*11 RNAi-treated nematodes compared to empty vector-treated nematodes (inserts sequenced up to 50nt).

**Supplementary Figure 2. Integrator localizes to sites of piRNA biogenesis in germ cells (relates to Figure 2)**.

A. Quantification of the number of INTS-6::GFP and mCherry::PRDE-1 foci/cell and their co-localization across the *C. elegans* germline. 1-5 correspond to the germline sections shown in Figure 2B.

B. Co-localization of FLAG::INTS-11 and mCherry::PRDE-1 in the *C. elegans* germline observed by anti-FLAG and anti-mCherry immunofluorescent staining.

**Supplementary Figure 3. piRNA precursor length distribution of individual replicates (relates to Figure 3)**.

A. Boxplots comparing the length of pairs of nucleoplasmic and chromatin-libraries corresponding to the same initial nuclei sample.

B. Length distribution of chromatin-bound and nucleoplasmic piRNA precursors in the two individual replicates of corresponding to each condition.

**Supplementary Figure 4. Further analysis of cleavage fragment abundance (relates to Figure 4)**.

A. Heatmap showing the coverage of unique sequences mapping to piRNA loci after removal of reads corresponding to mature 21U-RNAs (see Methods). The first peak corresponds to 21U-RNA degradation products and 5′ PPP initiation products starting at piRNA TSSs. The second peak corresponds to degradation fragments initiating at +38 on average (Figure 4D), resulting in sequence coverage peaking at +48 due to a median length of fragments of 20 nt (Figure 4D).

B. Log_2_ fold change distributions of degradation fragment counts per million of mapped reads relative to N2 EV. Fold changes were calculated after addition of 0.01 pseudocpms. Two independent biological replicates are shown.

C. Positions of 3′ ends of 5′ P fragments relative to piRNA TSSs. Signal is normalized to counts per million mapped reads.

D. Number of loci with detectable cleavage fragments across bins of loci stratified according to the strength of their termination signals, from weak (GC-rich) to strong (AT-rich). Data from each individual library is shown.

**Supplementary Figure 5. Integrator and the strength of AT-rich termination signals (relates to Figure 4)**.

Length distribution of chromatin-bound motif-dependent piRNA precursors stratified by the strength of downstream termination signals from weak (GC-rich) to strong (AT-rich), in empty vector and *ints-*11 RNAi-treated nematodes. Two replicates for each condition are shown.

**Supplementary Figure 6. Nascent SLIC-CAGE detects readthrough transcription from piRNA loci (relates to Figure 5)**.

A. Clustered correlation matrix of promoter-mapping CAGE tpms for the top 1000 genes with the highest CAGE signal.

B. Intron retention ratio distributions across nascent RNA samples, and in a total RNA control, estimated through IRFinder analysis.

C-D. Total raw CAGE tags mapping between −10 and −1 nucleotide positions upstream of motif-dependent piRNAs (C) and motif-independent piRNAs (D).

E. Total unique CAGE-detected TSSs mapping between −10 and −1 nucleotide positions upstream of motif-independent piRNAs.

F. Mean normalized expression levels of CAGE-detected and non-detected motif-dependent piRNAs at the short-capped RNA level.

G. Downsampling analysis of CAGE libraries showing that the detection of piRNA loci is saturated with sequencing depth. Total detected loci in each of the two replicates were averaged for each bootstrapped sample.

**Supplementary Figure 7. Accessibility and H3K27 acetylation levels at piRNA promoters (relates to Figure 5)**.

A. ATAC-seq read coverage around motif-dependent and motif-independent piRNA promoters, in wild-type young adult wild-type worms, and in day 1 adult germlineless glp-1 mutant worms lacking a germline. ‘Motif-dependent CAGE’ refers to motif-dependent loci detected by CAGE.

B. Log_2_ ratio in accessibility signal at motif-dependent piRNA loci between young adult wild-type worms and day 1 adult germlineless glp-1 mutants lacking a germline. This reflects a specific increase in accessibility at motif-dependent loci in germ cells. ‘Motif-dependent CAGE’ refers to motif-dependent loci detected by CAGE.

C. H3K27ac ChIP signal minus input around motif-dependent and motif-independent piRNA loci in isolated germ nuclei (left panel), and in isolated somatic nuclei (right panel).

D-E. Distributions of H3K27ac (D) and accessibility signal (E) in motif-dependent and motif-independent piRNA loci.

**Supplementary Figure 8. Transcriptional changes in snRNAs and protein-coding genes upon *ints-*11 knockdown (relates to Figure 5)**.

A. Nascent CAGE signal at snRNA genes upon *ints-*11 knockdown, in a wild-type and a *tfiis* mutant background. Extensive readthrough transcription with ∼500nt fragments is readily detectable upon *ints-*11 knockdown.

B. Distributions of log_2_ fold changes in nascent CAGE signal at snRNA loci upon *ints-*11 knockdown, in a wild-type and a *tfiis* mutant background, using N2 empty vector as a baseline.

C. MAplot showing differentially expressed protein-coding genes at the nascent short capped RNA level, with an excess of upregulated genes.

D. MAplot showing differentially expressed protein-coding genes detected by nascent CAGE, with an excess of upregulated genes.

E. Overlap between upregulated genes in the nascent scRNA and CAGE datasets.

F. Nascent CAGE signal at the set of upregulated protein coding genes upon *ints-*11 knockdown, in a wild-type and a *tfiis* mutant background.

G. Correlation between changes in nascent short-capped RNA and CAGE signal, for genes detected as differentially expressed in at least one of the two assays.

H. Log_2_ odds ratios of enrichment of motif-independent piRNA loci in protein-coding gene promoters detected as differentially expressed upon *ints-*11 knockdown, at the nascent scRNA level, at the CAGE level, or both (see labels).

## Supplementary tables

**Table S1.** *C. elegans* strains.

| Strain | Genotype | Description | Reference |
| --- | --- | --- | --- |
| SX1316 | mjls144 II | piRNA sensor strain | (Bagijn et al., 2012b) |
| JCP383 | ints-6 (tm1615) IV; jcpSi10 [pJC51 (ints-6p::ints-6::3xFLAG::eGFP::ints-6UTR, unc-119(+)) II | INTS-6::eGFP rescue strain | (Gómez-Orte et al., 2019) |
| SX2700 | mjSi74 I; mjIs144 II; prde-1(mj207) | mCherry::PRDE-1 rescue strain | (Weick et al., 2014) |
| TEE172 | ints-6 (tm1615) IV; jcpSi10 [pJC51 (ints-6p::ints-6::3xFLAG::eGFP::ints-6UTR, unc-119(+)) II; mjSi74 I | Generated by crossing JCP383 and SX2700 | This study |

**Table S2.**
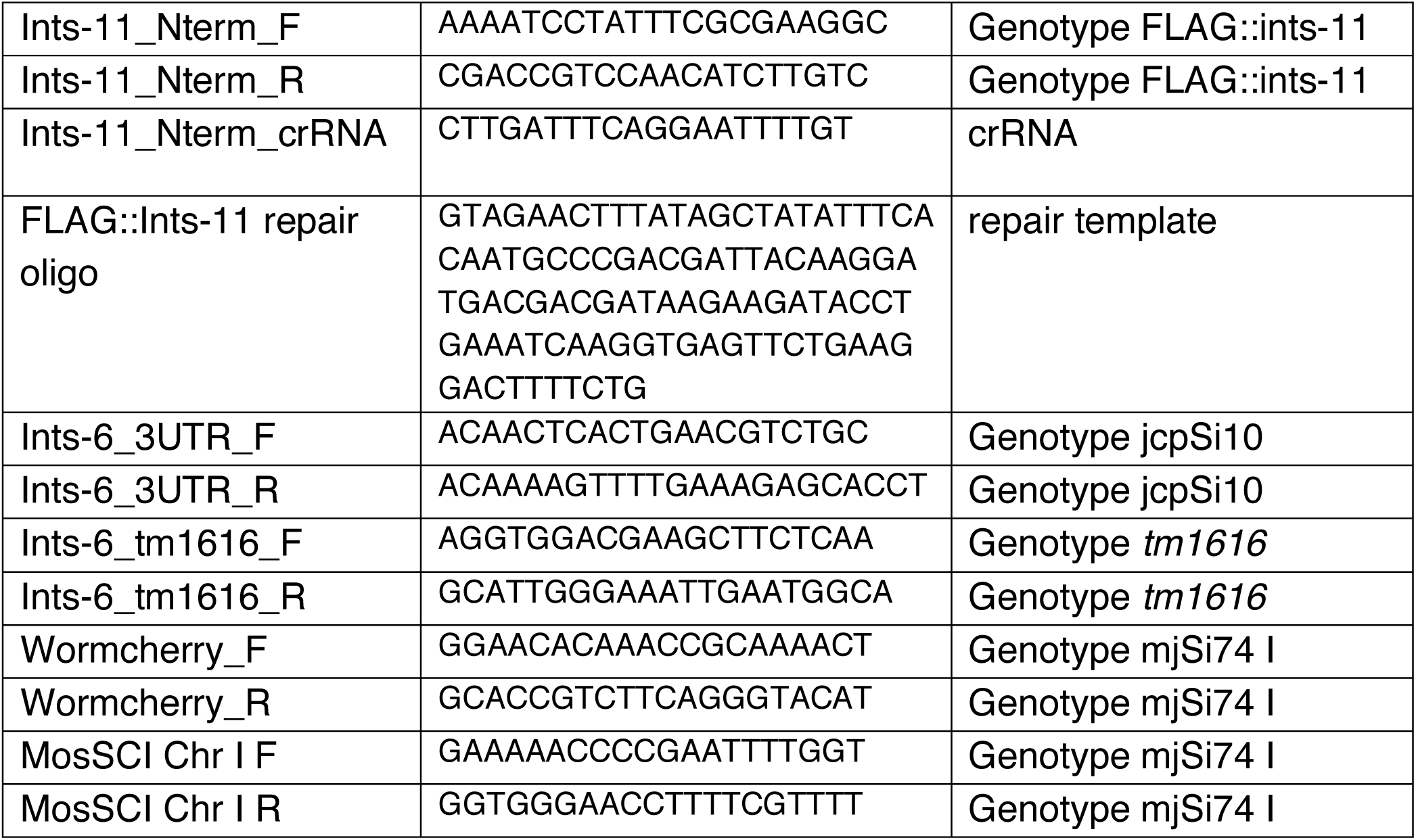
Oligonucleotides.

## References

Aoi, Y., Smith, E.R., Shah, A.P., Rendleman, E.J., Marshall, S.A., Woodfin, A.R., Chen, F.X., Shiekhattar, R., and Shilatifard, A. (2020). NELF Regulates a Promoter-Proximal Step Distinct from RNA Pol II Pause-Release. Mol. Cell.

Awrey, D.E., Weilbaecher, R., Hemming, S.A., Orlicky, S.M., Kane, C., and Edwards, A.M. (1997). Transcription elongation through DNA arrest sites - A multistep process involving both RNA polymerase II subunit RPB9 and TFIIS. J. Biol. Chem. 272, 14747–14754.

Bagijn, M.P., Goldstein, L.D., Sapetschnig, A., Weick, E.-M., Bouasker, S., Lehrbach, N.J., Simard, M.J., Miska, E.A., Das, P.P., Batista, P.J., et al. (2012a). Function, targets, and evolution of Caenorhabditis elegans piRNAs. Science 337, 574–578.

Bagijn, M.P., Goldstein, L.D., Sapetschnig, A., Weick, E., Bouasker, S., Lehrbach, N.J., Simard, M.J., and Miska, E. a (2012b). Function, Targets and Evolution of Caenorhabditis elegans piRNAs. Science (80-.). 574.

Baillat, D., Hakimi, M.-A., Näär, A.M., Shilatifard, A., Cooch, N., and Shiekhattar, R. (2005). Integrator, a Multiprotein Mediator of Small Nuclear RNA Processing, Associates with the C-Terminal Repeat of RNA Polymerase II. Cell 123, 265–276.

Balwierz, P.J., Carninci, P., Daub, C.O., Kawai, J., Hayashizaki, Y., Van Belle, W., Beisel, C., and van Nimwegen, E. (2009). Methods for analyzing deep sequencing expression data: constructing the human and mouse promoterome with deepCAGE data. Genome Biol. 10, R79.

Batista, P.J., Ruby, J.G., Claycomb, J.M., Chiang, R., Fahlgren, N., Kasschau, K.D., Chaves, D.A., Gu, W., Vasale, J.J., Duan, S., et al. (2008). PRG-1 and 21U-RNAs Interact to Form the piRNA Complex Required for Fertility in C. elegans. Mol. Cell 31, 67–78.

Beltran, T., Barroso, C., Birkle, T.Y., Stevens, L., Schwartz, H.T., Sternberg, P.W., Fradin, H., Gunsalus, K., Piano, F., Sharma, G., et al. (2019). Comparative Epigenomics Reveals that RNA Polymerase II Pausing and Chromatin Domain Organization Control Nematode piRNA Biogenesis. Dev. Cell 48, 793–810.e6.

Billi, A.C., Freeberg, M.A., Day, A.M., Chun, S.Y., Khivansara, V., and Kim, J.K. (2013). A Conserved Upstream Motif Orchestrates Autonomous, Germline-Enriched Expression of Caenorhabditis elegans piRNAs. PLoS Genet. 9.

Brennecke, J., Aravin, A.A., Stark, A., Dus, M., Kellis, M., Sachidanandam, R., and Hannon, G.J. (2007). Discrete Small RNA-Generating Loci as Master Regulators of Transposon Activity in Drosophila. Cell 128, 1089–1103.

Cecere, G., Zheng, G.X.Y., Mansisidor, A.R., Klymko, K.E., and Grishok, A. (2012). Promoters Recognized by Forkhead Proteins Exist for Individual 21U-RNAs. Mol. Cell 47, 734–745.

Chen, R.A.-J., Down, T.A., Stempor, P., Chen, Q.B., Egelhofer, T.A., Hillier, L.W., Jeffers, T.E., and Ahringer, J. (2013). The landscape of RNA polymerase II transcription initiation in C. elegans reveals promoter and enhancer architectures. Genome Res. 23, 1339–1347.

Cordeiro Rodrigues, R.J., de Jesus Domingues, A.M., Hellmann, S., Dietz, S., de Albuquerque, B.F.M., Renz, C., Ulrich, H.D., Sarkies, P., Butter, F., and Ketting, R.F. (2019). PETISCO is a novel protein complex required for 21U RNA biogenesis and embryonic viability. Genes Dev. 33, 857–870.

Creyghton, M.P., Cheng, A.W., Welstead, G.G., Kooistra, T., Carey, B.W., Steine, E.J., Hanna, J., Lodato, M.A., Frampton, G.M., Sharp, P.A., et al. (2010). Histone H3K27ac separates active from poised enhancers and predicts developmental state. Proc. Natl. Acad. Sci. 107, 21931 LP–21936.

Cvetesic, N., Leitch, H.G., Borkowska, M., Müller, F., Carninci, P., Hajkova, P., and Lenhard, B. (2018). SLIC-CAGE: high-resolution transcription start site mapping using nanogram-levels of total RNA. Genome Res.

Cvetesic, N., Pahita, E., and Lenhard, B. (2019). Transcription Start Site Mapping Using Super-low Input Carrier-CAGE. JoVE e59805.

Das, P.P., Bagijn, M.P., Goldstein, L.D., Woolford, J.R., Lehrbach, N.J., Sapetschnig, A., Buhecha, H.R., Gilchrist, M.J., Howe, K.L., Stark, R., et al. (2008). Piwi and piRNAs Act Upstream of an Endogenous siRNA Pathway to Suppress Tc3 Transposon Mobility in the Caenorhabditis elegans Germline. Mol. Cell 31, 79–90.

Dobin, A., Davis, C.A., Schlesinger, F., Drenkow, J., Zaleski, C., Jha, S., Batut, P., Chaisson, M., and Gingeras, T.R. (2013). STAR: ultrafast universal RNA-seq aligner. Bioinformatics 29, 15–21.

Dong, L., Cornaglia, M., Krishnamani, G., Zhang, J., Mouchiroud, L., Lehnert, T., Auwerx, J., and Gijs, M.A.M. (2018). Reversible and long-term immobilization in a hydrogel-microbead matrix for high-resolution imaging of Caenorhabditis elegans and other small organisms. PLoS One 13, e0193989.

Egloff, S., O’Reilly, D., Chapman, R.D., Taylor, A., Tanzhaus, K., Pitts, L., Eick, D., and Murphy, S. (2007). Serine-7 of the RNA Polymerase II CTD Is Specifically Required for snRNA Gene Expression. Science (80-.). 318, 1777–1779.

Egloff, S., Szczepaniak, S.A., Dienstbier, M., Taylor, A., Knight, S., and Murphy, S. (2010). The integrator complex recognizes a new double mark on the RNA polymerase II carboxyl-terminal domain. J. Biol. Chem. 285, 20564–20569.

Elrod, N.D., Henriques, T., Huang, K.-L., Tatomer, D.C., Wilusz, J.E., Wagner, E.J., and Adelman, K. (2019). The Integrator Complex Attenuates Promoter-Proximal Transcription at Protein-Coding Genes. Mol. Cell 76, 738–752.e7.

Frey, M.R., and Matera, A.G. (2001). RNA-mediated interaction of Cajal bodies and U2 snRNA genes. J. Cell Biol. 154, 499 LP–510.

Gardini, A., Baillat, D., Cesaroni, M., Hu, D., Marinis, J.M., Wagner, E.J., Lazar, M.A., Shilatifard, A., and Shiekhattar, R. (2014). Integrator Regulates Transcriptional Initiation and Pause Release following Activation. Mol. Cell 56, 128–139.

Gómez-Orte, E., Sáenz-Narciso, B., Zheleva, A., Ezcurra, B., de Toro, M., López, R., Gastaca, I., Nilsen, H., Sacristán, M.P., Schnabel, R., et al. (2019). Disruption of the Caenorhabditis elegans Integrator complex triggers a non-conventional transcriptional mechanism beyond snRNA genes. PLOS Genet. 15, e1007981.

Gu, W., Lee, H.-C., Chaves, D., Youngman, E.M., Pazour, G.J., Conte, D., and Mello, C.C. (2012). CapSeq and CIP-TAP identify Pol II start sites and reveal capped small RNAs as C. elegans piRNA precursors. Cell 151, 1488–1500.

Gunawardane, L.S., Saito, K., Nishida, K.M., Miyoshi, K., Kawamura, Y., Nagami, T., Siomi, H., and Siomi, M.C. (2007). A Slicer-Mediated Mechanism for Repeat-Associated siRNA 5’ End Formation in Drosophila. Science (80-.). 315, 1587 LP–1590.

Haberle, V., Forrest, A.R.R., Hayashizaki, Y., Carninci, P., and Lenhard, B. (2015). CAGEr: precise TSS data retrieval and high-resolution promoterome mining for integrative analyses. Nucleic Acids Res. 43, e51–e51.

Han, M., Wei, G., McManus, C.E., Hillier, L.W., and Reinke, V. (2019). Isolated C. elegans germ nuclei exhibit distinct genomic profiles of histone modification and gene expression. BMC Genomics 20, 500.

Handler, D., Meixner, K., Pizka, M., Lauss, K., Schmied, C., Gruber, F.S., and Brennecke, J. (2013). The genetic makeup of the Drosophila piRNA pathway. Mol. Cell 50, 762–777.

Jänes, J., Dong, Y., Schoof, M., Serizay, J., Appert, A., Cerrato, C., Woodbury, C., Chen, R., Gemma, C., Huang, N., et al. (2018). Chromatin accessibility dynamics across C. elegans development and ageing. Elife 7, e37344.

Kamath, R.S., and Ahringer, J. (2003). Genome-wide RNAi screening in Caenorhabditis elegans. Methods 30, 313–321.

Kasper, D.M., Wang, G., Gardner, K.E., Johnstone, T.G., and Reinke, V. (2014). The C.elegans SNAPc component SNPC-4 coats piRNA domains and is globally required for piRNA abundance. Dev. Cell.

Kozomara, A., and Griffiths-Jones, S. (2014). MiRBase: Annotating high confidence microRNAs using deep sequencing data. Nucleic Acids Res. 42, 68–73.

Lai, F., Gardini, A., Zhang, A., and Shiekhattar, R. (2015). Integrator mediates the biogenesis of enhancer RNAs. Nature 525, 399.

Langmead, B., and Salzberg, S.L. (2012). Fast gapped-read alignment with Bowtie 2. Nat Meth 9, 357–359.

Langmead, B., Trapnell, C., Pop, M., and Salzberg, S.L. (2009). Ultrafast and memory-efficient alignment of short DNA sequences to the human genome. Genome Biol. 10, R25.

Li, H., Handsaker, B., Wysoker, A., Fennell, T., Ruan, J., Homer, N., Marth, G., Abecasis, G., Durbin, R., and Subgroup, 1000 Genome Project Data Processing (2009). The Sequence Alignment/Map format and SAMtools. Bioinformatics 25, 2078–2079.

Maxwell, C.S., Kruesi, W.S., Core, L.J., Kurhanewicz, N., Waters, C.T., Lewarch, C.L., Antoshechkin, I., Lis, J.T., Meyer, B.J., and Baugh, L.R. (2014). Pol II Docking and Pausing at Growth and Stress Genes in C. elegans. Cell Rep. 6, 455–466.

Middleton, R., Gao, D., Thomas, A., Singh, B., Au, A., Wong, J.J.-L., Bomane, A., Cosson, B., Eyras, E., Rasko, J.E.J., et al. (2017). IRFinder: assessing the impact of intron retention on mammalian gene expression. Genome Biol. 18, 51.

Nojima, T., Gomes, T., Carmo-Fonseca, M., and Proudfoot, N.J. (2016). Mammalian NET-seq analysis defines nascent RNA profiles and associated RNA processing genome-wide. Nat. Protoc. 11, 413–428.

Nojima, T., Tellier, M., Foxwell, J., Ribeiro de Almeida, C., Tan-Wong, S.M., Dhir, S., Dujardin, G., Dhir, A., Murphy, S., and Proudfoot, N.J. (2018). Deregulated Expression of Mammalian lncRNA through Loss of SPT6 Induces R-Loop Formation, Replication Stress, and Cellular Senescence. Mol. Cell 72, 970–984.e7.

Ozata, D.M., Gainetdinov, I., Zoch, A., O’Carroll, D., and Zamore, P.D. (2019). PIWI-interacting RNAs: small RNAs with big functions. Nat. Rev. Genet. 20, 89–108.

Pritykin, Y., Brito, T., Schupbach, T., Singh, M., and Pane, A. (2017). Integrative analysis unveils new functions for the Drosophila Cutoff protein in noncoding RNA biogenesis and gene regulation. RNA 23, 1097–1109.

Quinlan, A.R., and Hall, I.M. (2010). BEDTools: a flexible suite of utilities for comparing genomic features. Bioinformatics 26, 841–842.

Ramírez, F., Ryan, D.P., Grüning, B., Bhardwaj, V., Kilpert, F., Richter, A.S., Heyne, S., Dündar, F., and Manke, T. (2016). deepTools2: a next generation web server for deep-sequencing data analysis. Nucleic Acids Res. 44, W160–W165.

Ruby, J.G., Jan, C., Player, C., Axtell, M.J., Lee, W., Nusbaum, C., Ge, H., and Bartel, D.P. (2006). Large-Scale Sequencing Reveals 21U-RNAs and Additional MicroRNAs and Endogenous siRNAs in C. elegans. Cell 127, 1193–1207.

Sigurdsson, S., Dirac-Svejstrup, A.B., and Svejstrup, J.Q. (2010). Evidence that transcript cleavage is essential for RNA polymerase II transcription and cell viability. Mol. Cell 38, 202–210.

Stadelmayer, B., Micas, G., Gamot, A., Martin, P., Malirat, N., Koval, S., Raffel, R., Sobhian, B., Severac, D., Rialle, S., et al. (2014). Integrator complex regulates NELF-mediated RNA polymerase II pause/release and processivity at coding genes. Nat. Commun. 5, 5531.

Wang, Q., Sawyer, I.A., Sung, M.-H., Sturgill, D., Shevtsov, S.P., Pegoraro, G., Hakim, O., Baek, S., Hager, G.L., and Dundr, M. (2016). Cajal bodies are linked to genome conformation. Nat. Commun. 7, 10966.

Weick, E.M., Sarkies, P., Silva, N., Chen, R. a., Moss, S.M.M., Cording, A.C., Ahringer, J., Martinez-Perez, E., and Miska, E. a. (2014). PRDE-1 is a nuclear factor essential for the biogenesis of Ruby motif-dependent piRNAs in C. elegans. Genes Dev. 28, 783–796.

Weng, C., Kosalka, J., Berkyurek, A.C., Stempor, P., Feng, X., Mao, H., Zeng, C., Li, W.-J., Yan, Y.-H., Dong, M.-Q., et al. (2019). The USTC co-opts an ancient machinery to drive piRNA transcription in C. elegans. Genes Dev. 33, 90–102.

Zeng, C., Weng, C., Wang, X., Yan, Y.-H., Li, W.-J., Xu, D., Hong, M., Liao, S., Dong, M.-Q., Feng, X., et al. (2019). Functional Proteomics Identifies a PICS Complex Required for piRNA Maturation and Chromosome Segregation. Cell Rep. 27, 3561–3572.e3.

