## Supplementary figures and images for "Integrator terminates promoter-proximal Pol II to generate *C. elegans* piRNA precursors"

### Supplemental Figures

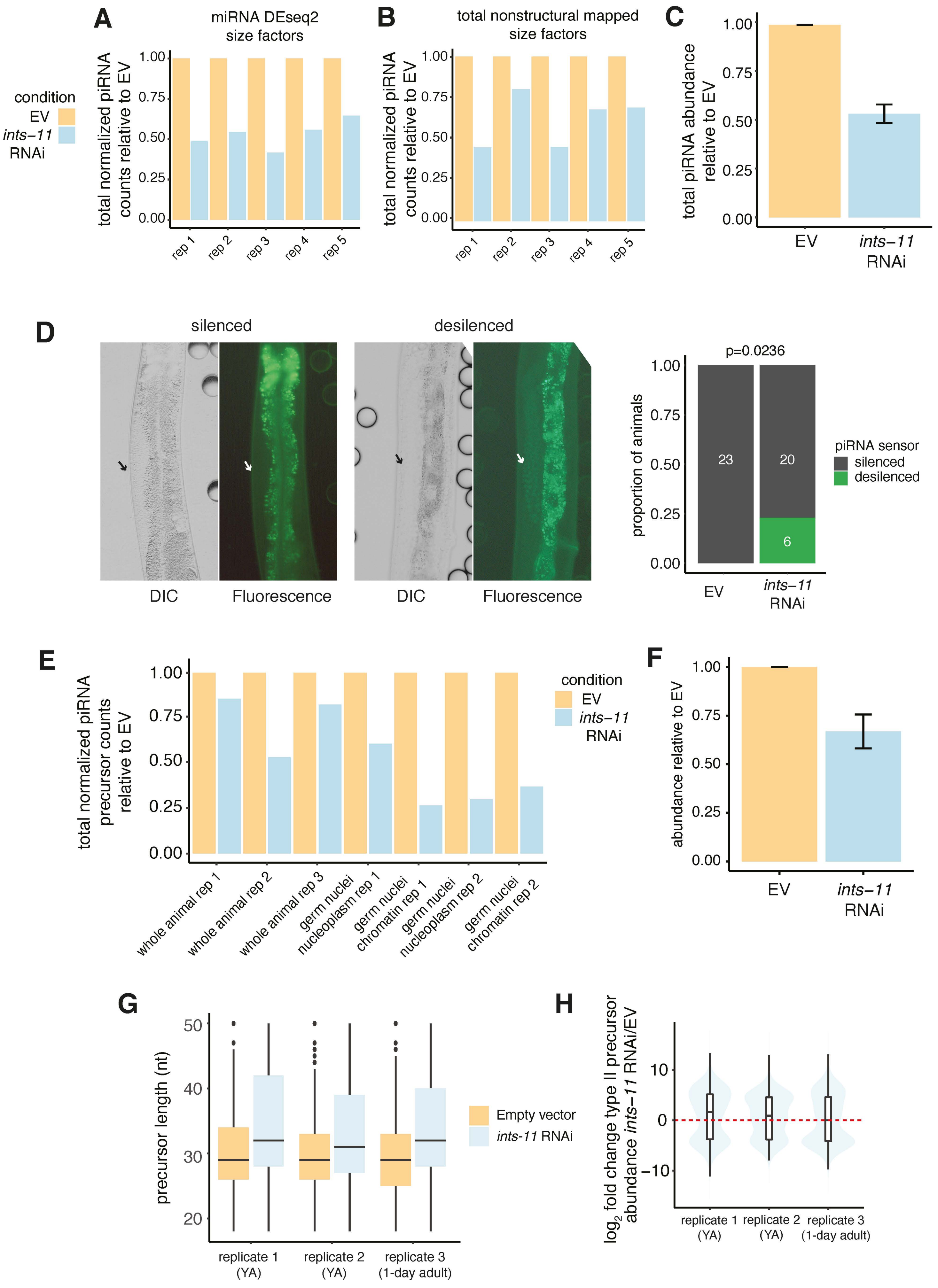

**A**

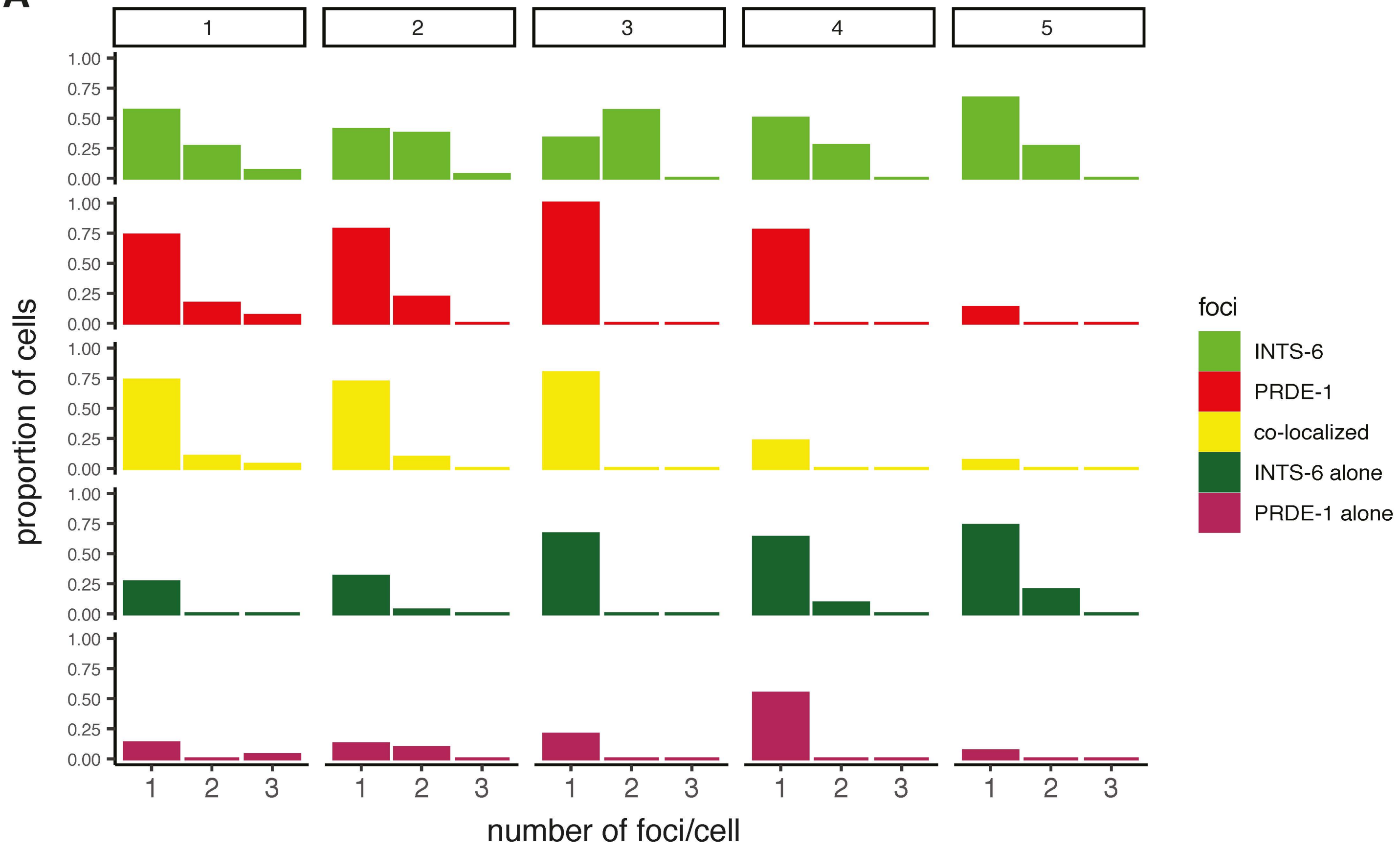


**B**

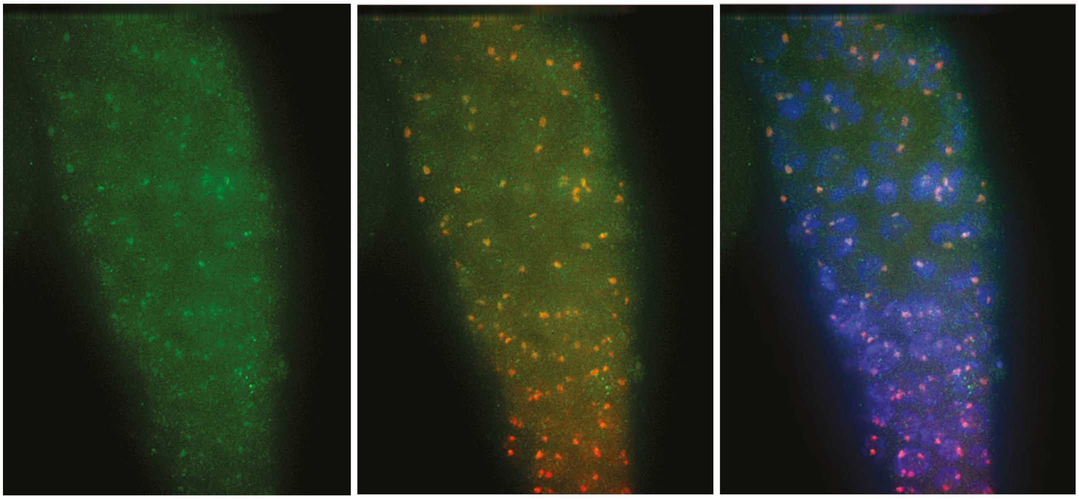

FLAG::ints-11

FLAG::ints-11
mCherry::PRDE-1

FLAG::ints-11
mCherry::PRDE-1
DAPI

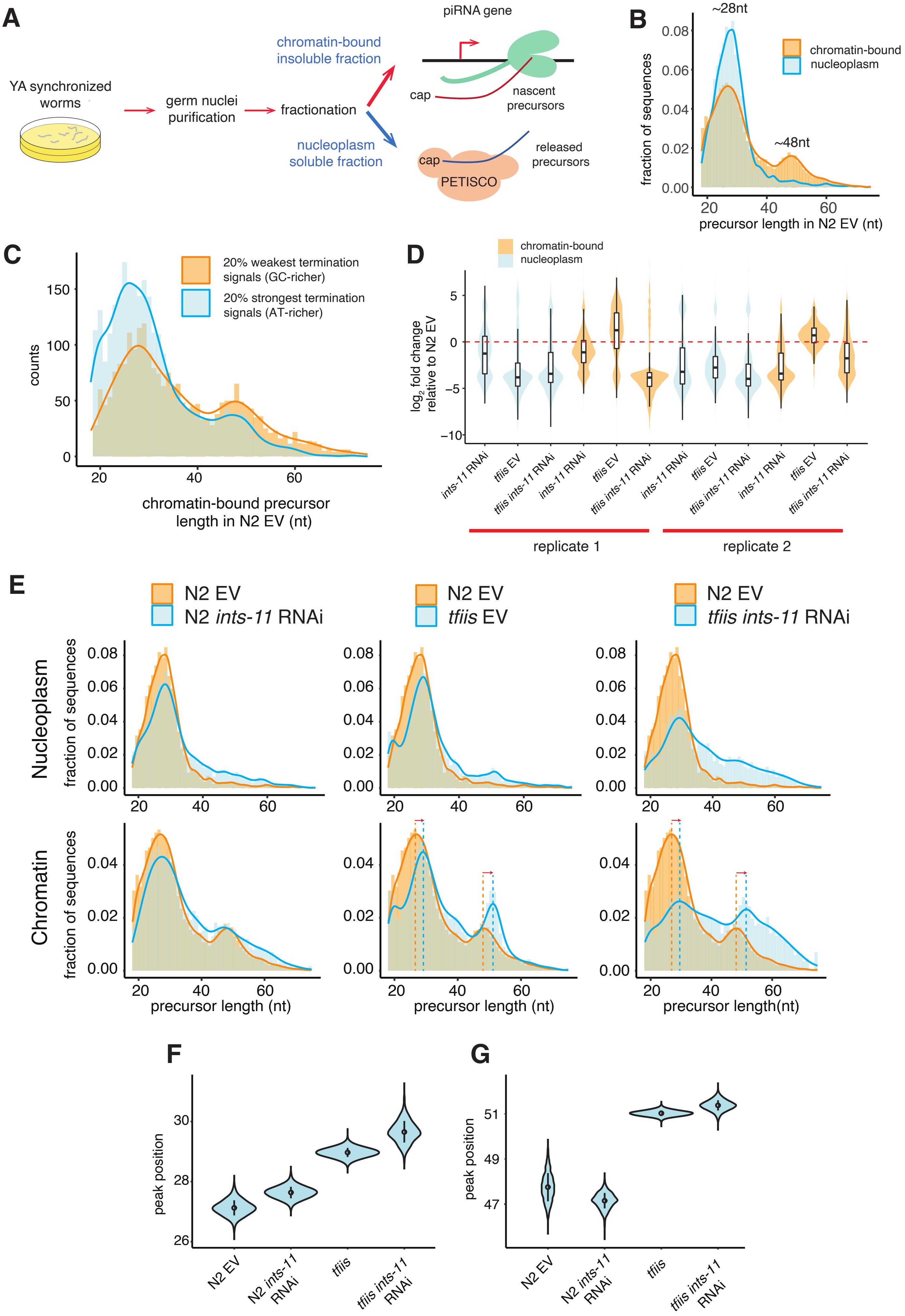

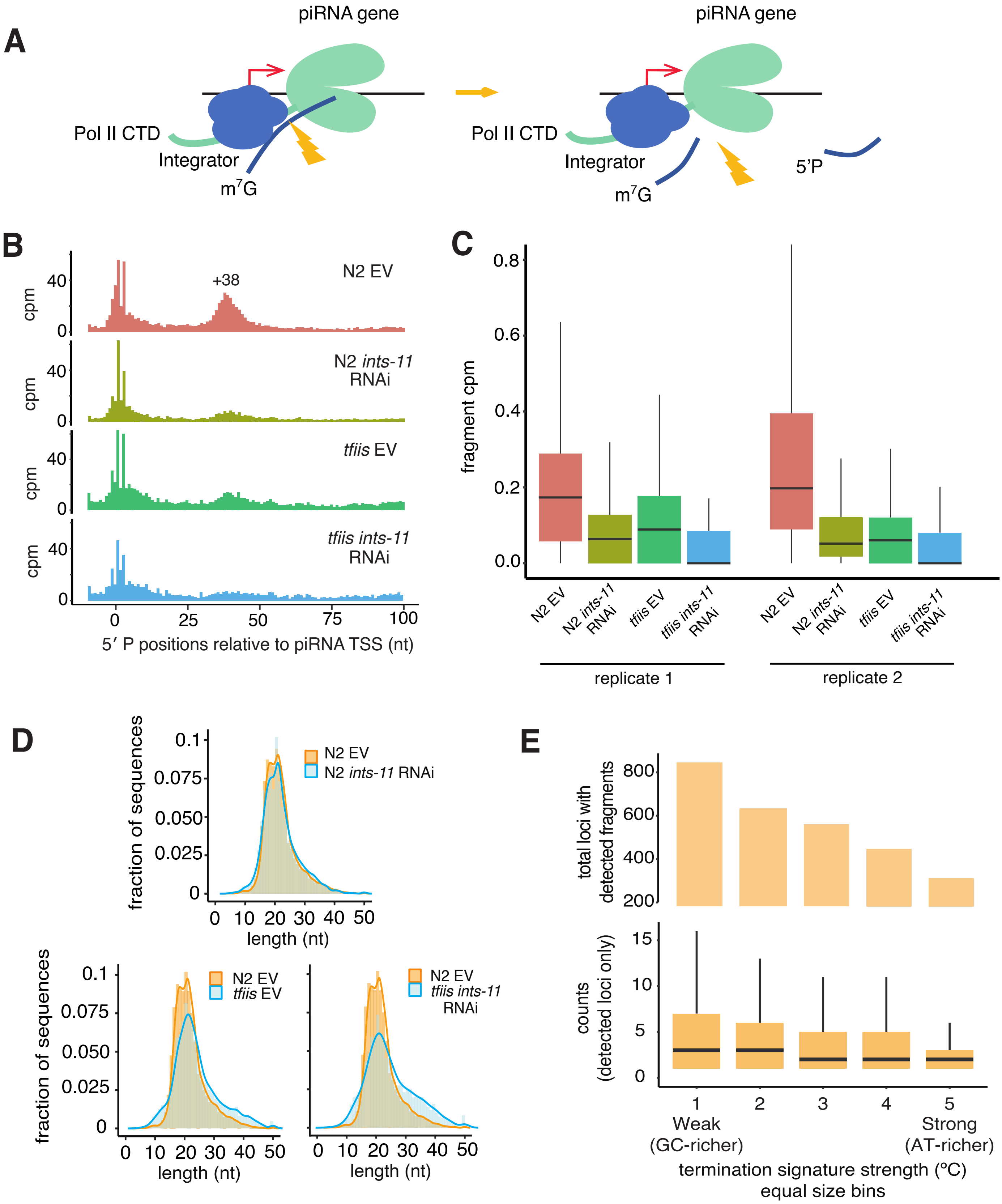

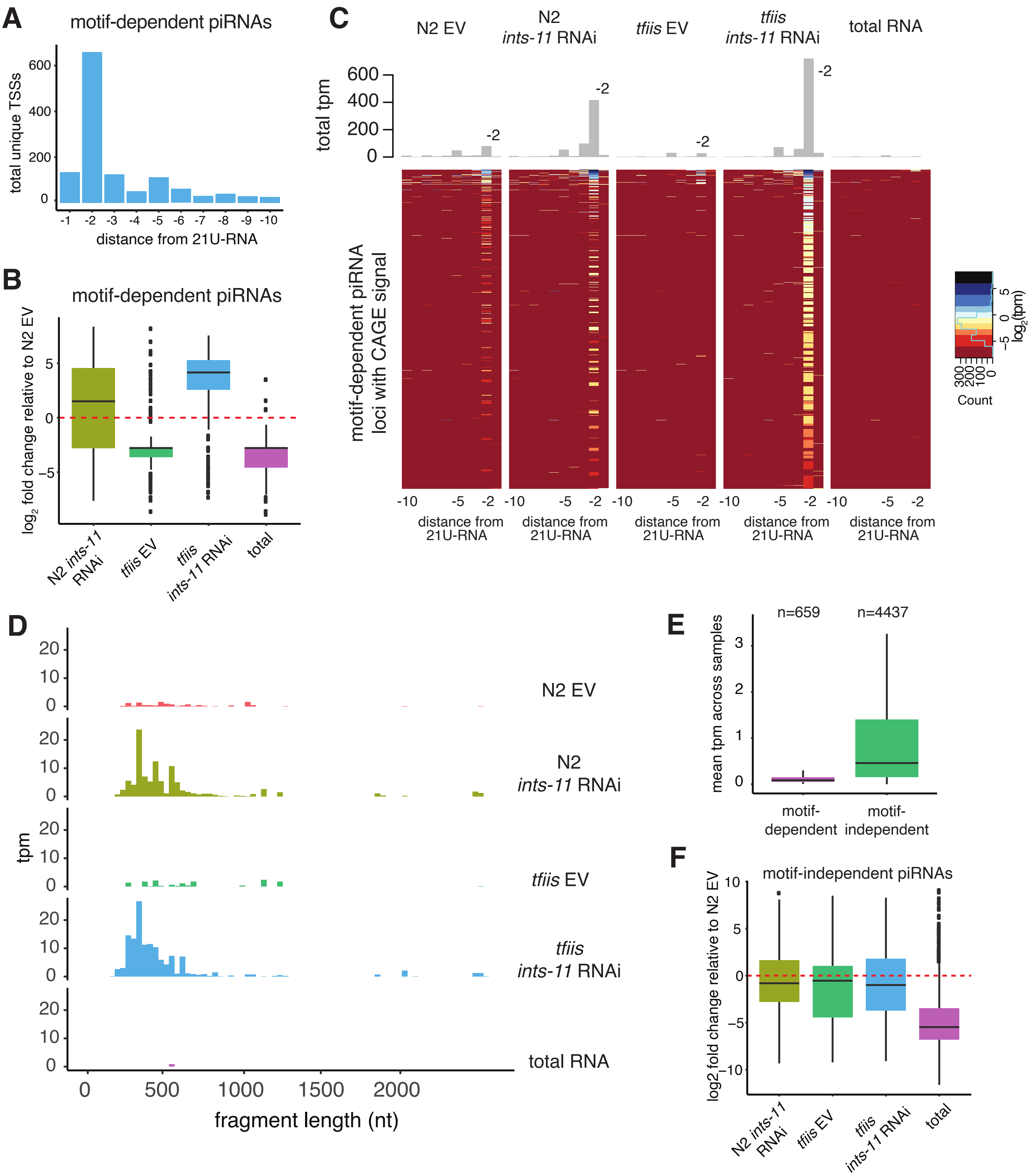

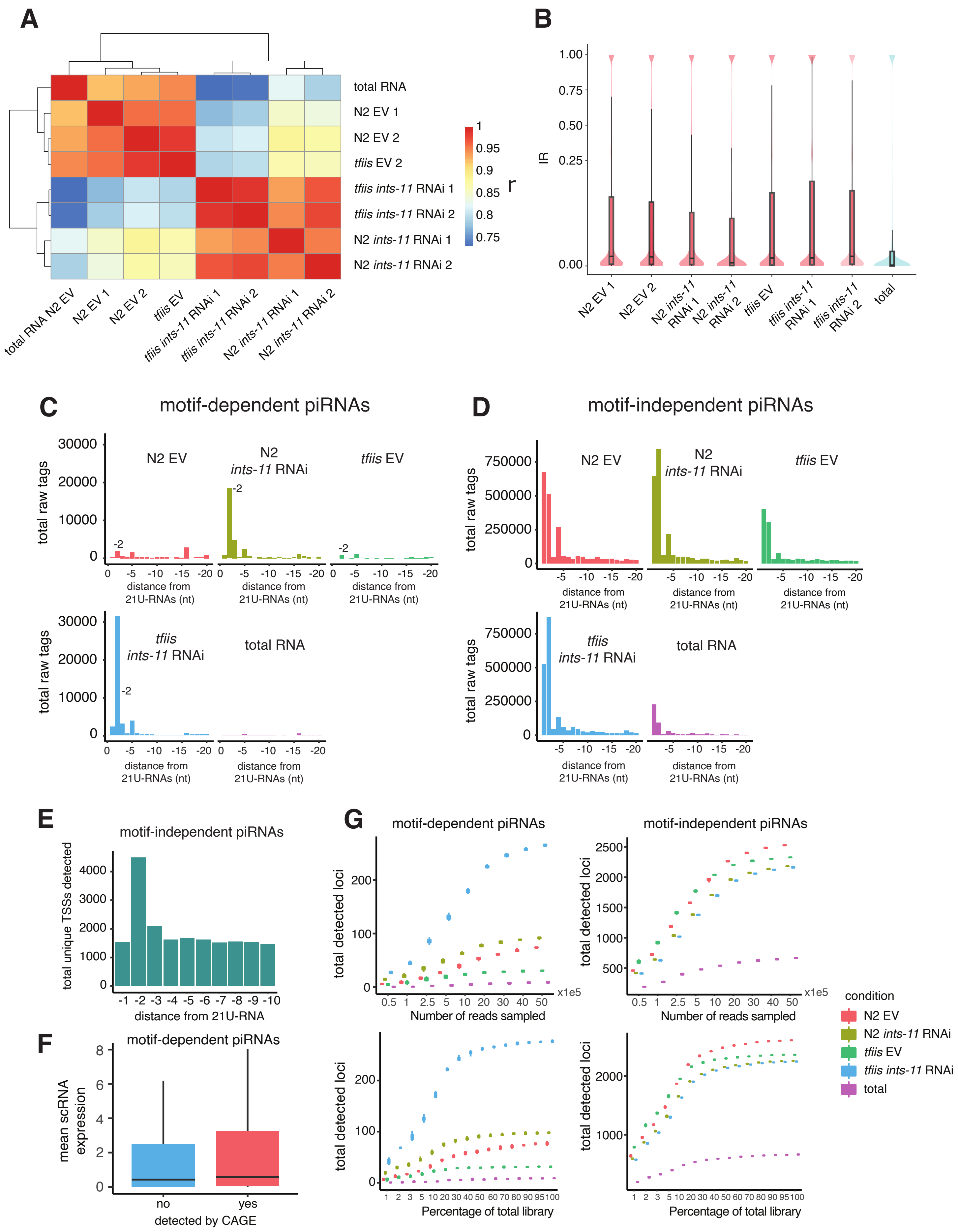

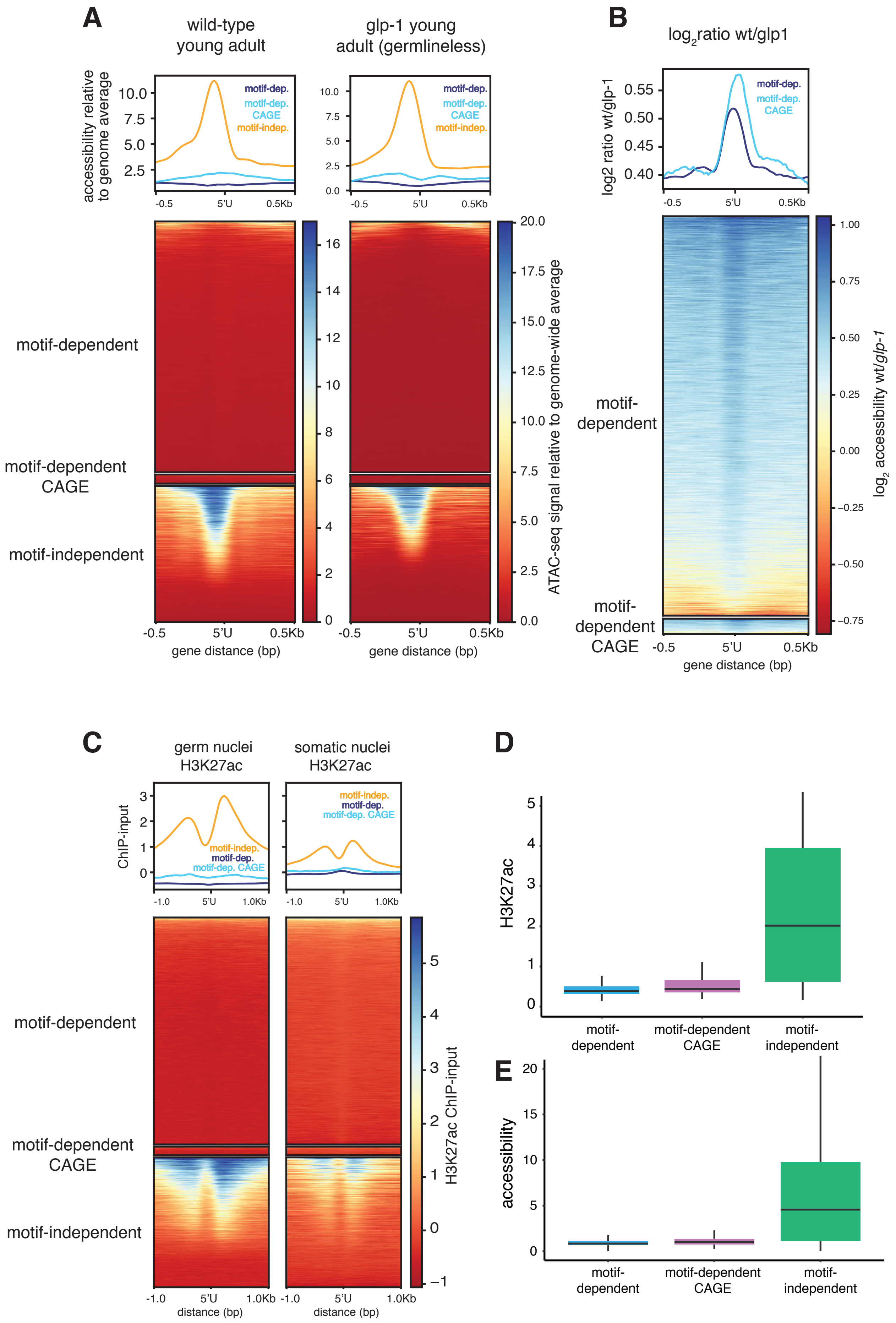

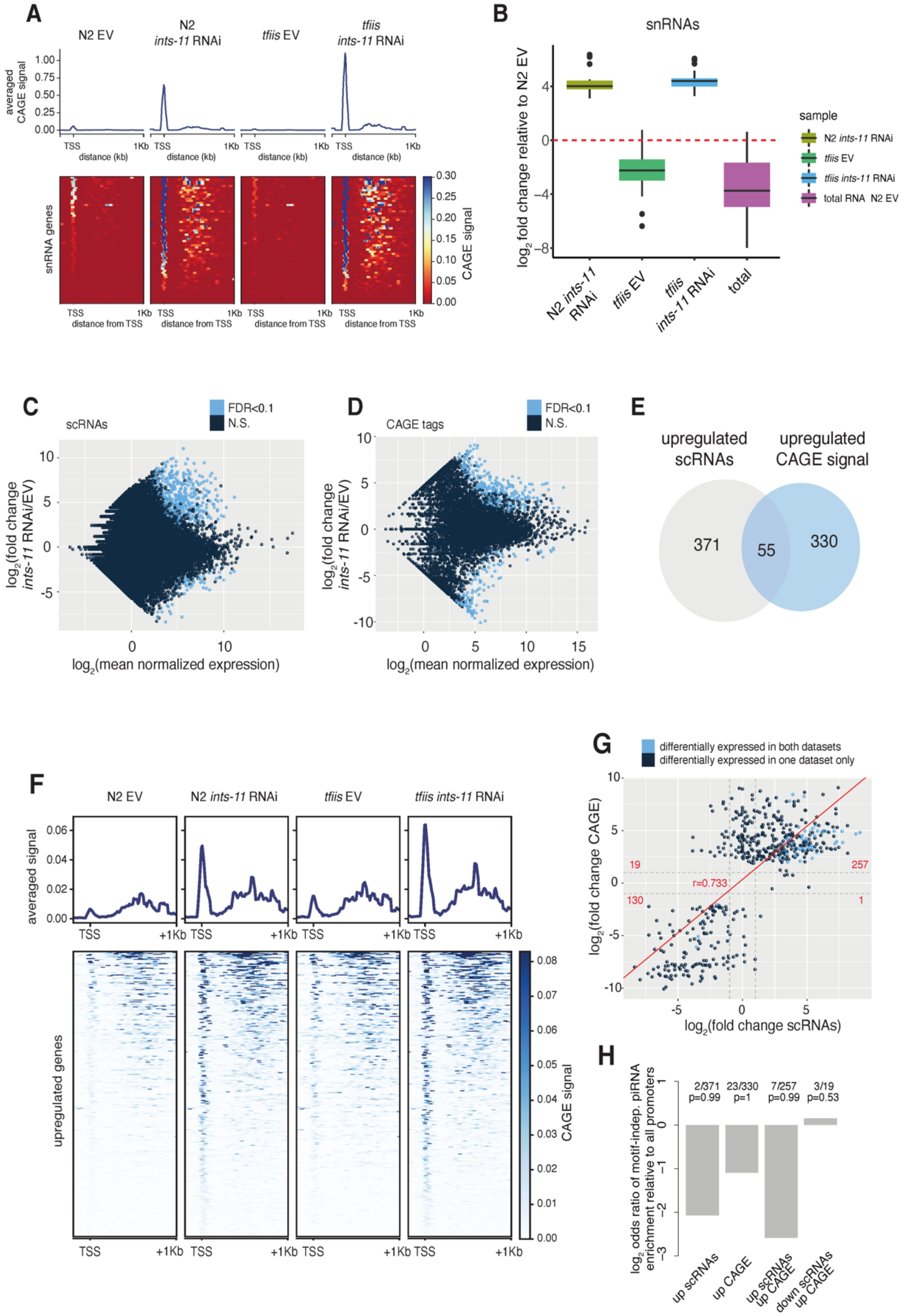
